# Genomic heritability of song and condition in wild singing mice

**DOI:** 10.1101/2020.10.08.321141

**Authors:** Tracy T. Burkhard, Mikhail Matz, Steven M. Phelps

## Abstract

Courtship displays are dramatic examples of complex behaviors that vary within and among species. Evolutionary explanations for this diversity rely upon genetic variation, yet the heritability of complex phenotypes is seldom investigated in the field. Here, we estimate genomic heritability of advertisement song and body condition in a wild population of singing mice. The heritability of song exhibits a systematic pattern, with high heritability for spectral characteristics linked to vocal morphology, intermediate heritability for rhythmic patterns, and lower but significant heritability for measures of motivation, like song length and rate. Physiological measures of condition, like hormonal markers of adiposity, exhibited intermediate heritability. Among singing mice, song rate and body condition have a strong phenotypic correlation; our estimate suggests a comparable genetic correlation that merits further study. Our results illustrate how advances in genomics and quantitative genetics can be integrated in free-living species to address longstanding challenges in behavior and evolution.

## Introduction

A fundamental challenge in evolutionary biology is to understand the origins of phenotypic diversity. The concept of heritability (*h*^*2*^)—the proportion of phenotypic variance due to genetic differences in a population (Falconer, 1981; Visscher et al., 2008)—is crucial to this understanding because it predicts a trait’s evolutionary response to selection (Falconer, 1981; Fisher, 1930; Lande, 1976; Price et al., 1991; Visscher et al., 2008). Traits of high heritability are more responsive to selection than traits of low heritability (Falconer, 1981; Lande, 1976), knowledge long exploited by livestock and agriculture breeders (Cassell et al., 2009; Dudley and Lambert, 2010; Hill and Kirkpatrick, 2010; Visscher et al., 2008) and famously illustrated by the rapid, repeated adaptation of highly heritable beak size in Darwin’s finches (Grant and Grant, 1993, 1995).

Few traits pose such interesting and challenging examples of phenotypic evolution as male sexual signals. Advertisement displays are present across diverse taxonomic groups (Wiens and Tuschhoff, 2020), vary dramatically within and among species (within: Dubuc et al., 2014; Izzo and Tibbetts, 2015; Karin et al., 2018; between: Ingram et al., 2016; Vanhooydonck et al., 2009), and evolve under both natural and sexual selection (Andersson, 1981; Endler, 1992). Heritable variation is not merely a substrate for this diversity but underlies common explanations for exaggerated male traits. Models of signal evolution by female choice assume heritable variation of male display, for example, but a popular subset also predicts genetic correlations between sexual display and male fitness or condition—correlations that are interpreted to indicate that females use displays to gauge the heritable fitness of males (Andersson, 1986; Hamilton and Zuk, 1982; Kirkpatrick and Ryan, 1991; Rowe and Houle, 1996).

Of course, not all variation in displays is heritable: sexual displays are complex, encompassing both stable structural elements and the dynamic behaviors that animate them. Relative to stable morphological elements of display, behavioral components are often highly specific to social and environmental contexts. For example, animals increase signaling effort (*e.g.* display rate or duration) in the presence of an audience, a widespread phenomenon that has been documented in vocalizing hyraxes (Demartsev et al., 2014), courting betta fish (Doutrelant et al., 2001), and many other taxa (reviewed in Zuberbühler, 2008). Behavioral components may also rely on male body condition (Cotton et al., 2004; Johnstone, 1993; Rowe and Houle, 1996), particularly if they are energetically expensive to produce (Chappell et al., 1995; Ophir et al., 2010) or incur the expense of fight or flight (Bachmann et al., 2017; Zuk and Kolluru, 1998). Different facets of display may thus reflect varying degrees of heritability. Even if selection pressures are consistent across these dimensions of display, sexual signals may evolve not as a single phenotype but as discrete components that undergo different rates of evolution (*e.g.* acoustic characteristics of frog calls, Cocroft and Ryan, 1995). Investigating the nature and extent of heritable variation underlying sexual signals is thus especially important to our broader understanding of how complex traits evolve.

Although heritability is essential to understanding evolution, it is challenging to study. By definition, heritability is highly population-specific – and its measures are sensitive to genetic variation and environmental circumstances (Falconer, 1981; Visscher et al., 2008; Waldmann, 2001; but see contrary evidence in Dochtermann et al., 2019; Waldmann, 2001). Despite its obvious value, our understanding of the quantitative genetics of wild populations has been limited due to the difficulties of ascertaining genetic relatedness in the field (Gienapp et al., 2017). One popular surrogate for heritability, especially in behavioral ecology, is repeatability, which eliminates the need for *a priori* knowledge of relatedness altogether (Bell et al., 2009; Boake, 1989; Dingemanse et al., 2002). Repeatability is facile and offers an upper bound estimate for heritability (Boake, 1989), but because direct extrapolation of repeatability to heritability can vary in reliability (*e.g.* can underestimate heritability, Dohm, 2002; St-Hilaire et al., 2017), its most appropriate use may be limited to hypothesis generation. Conversely, rigorous pedigree analysis, long the standard for quantitative genetics, is labor-intensive and impractical for many species that are either long-lived or difficult to track (Gienapp et al., 2017).

Recent advances in genomics and quantitative genetics afford new opportunities for assessing heritability in non-model and wild populations. In traditional pedigree analysis, pedigrees are fitted into animal models, linear (or generalized linear) mixed models first developed for animal breeding programs (Henderson, 1950; Hill and Kirkpatrick, 2010) and later applied to evolutionary questions (Kruuk, 2004; Lynch et al., 1998). The advent of reduced representation next-generation sequencing techniques, such as RAD-seq (Peterson et al., 2012), allows researchers to estimate population genome-wide relatedness without need for pedigree, marking a significant turning point for researchers interested in the quantitative genetics of wild populations. Genomic relatedness matrices (GRM) are estimated by identity-by-descent inference (Mousseau et al., 1998; Ritland, 1996) and can then be fitted into mixed models in lieu of pedigrees, resulting in estimates of “SNP-based” or “genomic” heritability. Researchers in human genetics, biostatistics, and quantitative genetics have developed computational tools that accommodate complex genomic models, which has led to important insights into complex human traits (*e.g.* Yang et al., 2010). Researchers in ecology and evolution have recently begun to apply these methods in non-model species to investigate heritable variation in the wild (Gervais et al., 2020, 2019; Perrier et al., 2018).

Here, we apply SNP-based methods to investigate patterns of heritability in singing behavior and body condition in a Costa Rican population of Alston’s singing mice (*Scotinomys teguina*). Singing mice are small, diurnal, and insectivorous muroid rodents of the family Cricetidae, eponymously named for their stereotyped advertisement songs (Hooper, 1972; Fig. 1). *Scotinomys* song makes a particularly interesting model for the study of signal evolution, being both acoustically complex and ecologically important. By the standards of muroid rodent vocalizations, songs are loud and long (7-10 s; Hooper and Carleton, 1976), comprising a rapidly repeated series of stereotyped, tonal frequency sweeps notes that span both audible nd ultrasonic (< 20kHz) frequencies. This suite of acoustic traits characterizes a notable departure from the short and simple ancestral ultrasonic vocalizations typical of most muroid rodents (Miller and Engstrom, 2007, 2012) and makes song detectable over long distances. Indeed, singing mouse song seems designed for that purpose, playing roles in both mate attraction and male-male competition (Fernández-Vargas et al., 2011; Hooper and Carleton, 1976; George, 2014; Pasch et al., 2011a, 2013). Interestingly, aspects of singing effort are predicted by male body condition (Burkhard et al., 2018; Pasch et al., 2011a), providing a putative mechanism for signaling decisions and female preferences.

**Figure 1:**
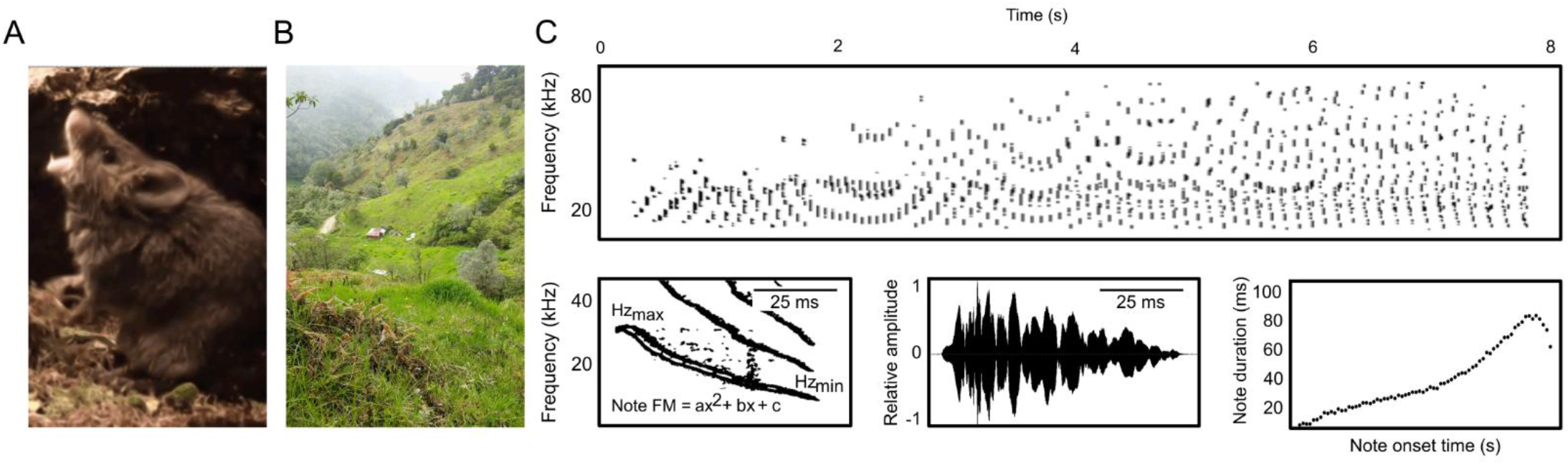
Singing behavior of a wild population of Alston’s singing mice (*Scotinomys teguina*). *(A)* Still from a video of a singing male mouse. *(B)* Typical grassland and forest habitat characterizing San Gerardo de Dota, Costa Rica. *(C)* Example spectrogram of an advertisement song. Lower panels: *left* and *middle* depict a close-up of a single note at 5 s during the song. *Left*, spectrogram detail of note with minimum and maximum frequency labeled. A quadratic curve was fit to each note to measure note frequency modulation (FM). FMa describes note curvature, FMb describes slope, and FMc describes starting frequency. *Middle*, waveform detail of note with peak amplitude, peak amplitude within first quarter of note, and note duration labeled. *Right*, note duration changes as a function of song length. Photo credit: *(A)* B. Pasch, *(B)* T. Burkhard.

To examine patterns of heritability within this species, we recorded and analyzed songs and singing behavior from 168 wild-caught mice from San Gerardo de Dota, Costa Rica (Fig. 1, Table 1), and we quantified acoustic repeatability within and variation among individuals. We measured a variety of condition phenotypes from these mice, including morphometric traits, plasma nutrients, and circulating hormone levels, and then examined the phenotypic relationships between condition and song (Fig. 2, Table 2). We generated genotype data for 157 mice using ddRAD-seq, and then deduced pairwise genetic covariance matrices. Finally, we used these data to fit animal models to explore patterns of heritability and co-heritability in song and condition phenotypes. Our study provides the first heritability estimates for complex signaling behaviors and their physiological substrates in a wild population and illustrates the utility of such methods to address evolutionary questions.

**Table 1:**
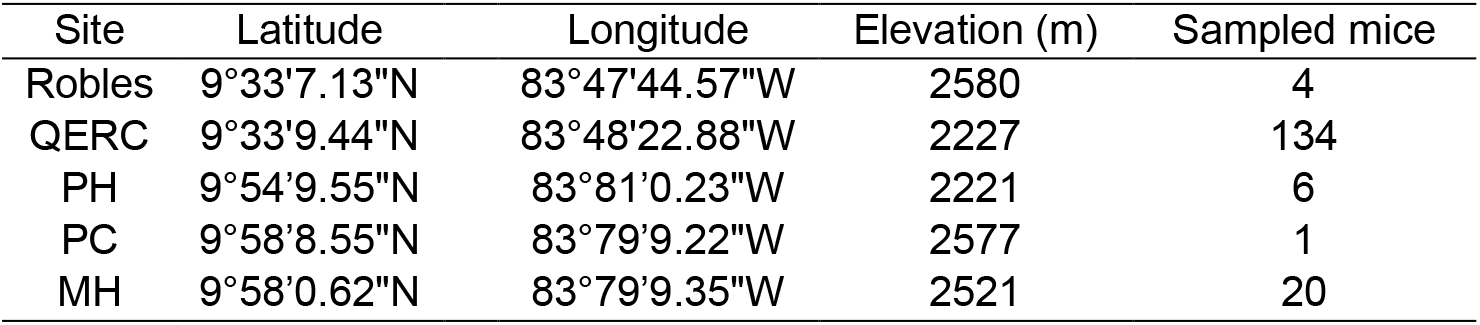
Trapping summary. Elevation taken at GPS waypoint.

**Table 2:** Sample sizes and summary statistics for each song and condition phenotype recorded from 165 singing mice.

| Phenotype | Units | N<sub>Total</sub> | N (♂) | N (♀) | Mean ± SD | Range |
| --- | --- | --- | --- | --- | --- | --- |
| Song rate | songs/2h | 129 | 97 | 32 | 2.1 ± 2.5 | 0 – 14 |
|  |  |  |  |  | 2.2 ± 2.3 (♂) | 0 – 9 (♂) |
|  |  |  |  |  | 1.8 ± 2.8 (♀) | 0 – 14 (♀) |
| Song length | s | 106 | 84 | 22 | 6.3 ± 1.5 | 4.1 – 8.8 |
|  |  |  |  |  | 6.7 ± 1.1 (♂) | 4.1 – 8.8 (♂) |
|  |  |  |  |  | 6.1 ± 0.9 (♀) | 4.4 – 7.7 (♀) |
| Trill rate | notes/s | 106 | 84 | 22 | 6.7 ± 1.1 | 8.5 – 15.9 |
|  |  |  |  |  | 11.8 ± 1.5 (♂) | 8.5 – 15.9 (♂) |
|  |  |  |  |  | 12.1 ± 1.1 (♀) | 9.1 – 13.6 (♀) |
| Note number | notes | 106 | 84 | 22 | 76.5 ± 12.8 | 43.5 – 104 |
|  |  |  |  |  | 77.5 ± 12.8 (♂) | 43.5 – 104 (♂) |
|  |  |  |  |  | 73.0 ± 12.6 (♀) | 55 – 104 (♀) |
| Dominant frequency | kHz | 106 | 84 | 22 | 24670.8 ± 3782.8 | 18052 – 35272 |
|  |  |  |  |  | 24596.1 ± 3856.7 (♂) | 18052 – 35272 (♂) |
|  |  |  |  |  | 24942.5 ± 3573.2 (♀) | 20079 – 32158 (♀) |
| Mean bandwidth | kHz | 106 | 84 | 22 | 25064.6 ± 7799.7 | 3306 – 45949 |
|  |  |  |  |  | 24439.3 ± 8209.9 (♂) | 3306 – 45949 (♂) |
|  |  |  |  |  | 27338.3 ± 5667.5 (♀) | 19599 – 44188 (♀) |
| Minimum frequency | kHz | 106 | 84 | 22 | 13660.62 ± 13856.53 | 15872 – 92570 |
|  |  |  |  |  | 14935.4 ± 15396.04 (♂) | 15872 – 92570 (♂) |
|  |  |  |  |  | 9025.06 ± 1740 (♀) | 36621 – 89264 (♀) |
| Maximum frequency | kHz | 106 | 84 | 22 | 55035.0 ± 18821.1 | 15872 – 92570 |
|  |  |  |  |  | 54309.9 ± 19123.3 (♂) | 15872 – 92570 (♂) |
|  |  |  |  |  | 57671.6 ± 17851.8 (♀) | 36621 – 89264 (♀) |
| Entropy | nat | 106 | 84 | 22 | -4.7 ± 0.4 | -5.6 – -4.0 |
|  |  |  |  |  | -4.6 ± 0.35 (♂) | -5.6 – -4.0 (♂) |
|  |  |  |  |  | -4.8 ± 0.4 (♀) | -5.3 – -4.0 (♀) |
| ddRAD | NA | 157 | 128 | 29 | NA |  |
| PC condition | NA | 52 | 52 | 0 |  |  |
| Mass | g | 165 | 133 | 32 | 12.6 ± 1.5 | 8 – 16.4 |
| Residual body mass | g | 165 | 133 | 32 | 0 ± 1 | -3.1 – 2.5 |
| Hindfoot | mm | 165 | 133 | 32 | 16.4 ± 0.6 | 15 – 18 |
| Anogenital distance | mm | 159 | 127 | 32 | 5.5 ± 1.3 | 2 – 8 |
|  |  |  |  |  | 2.2 ± 2.3 (♂) | 0 – 9 (♂) |
|  |  |  |  |  | 1.8 ± 2.8 (♀) | 0 – 14 (♀) |
| Glucose | ml dl<sup>-1</sup> | 78 | 78 | 0 | 111.9 ± 66.6 | 14.71 – 378.5 |
| Triglycerides | ml dl<sup>-1</sup> | 74 | 74 | 0 | 201.7 ± 119.5 | 52.2 – 718.3 |
| Adiponectin | ng ml<sup>-1</sup> | 63 | 63 | 0 | 132.8 ± 52.2 | 53.4 – 324.2 |
| Insulin | pg ml<sup>-1</sup> | 56 | 56 | 0 | 585.7 ± 502.7 | 69.0 – 2814.1 |
| Leptin | pg ml<sup>-1</sup> | 56 | 56 | 0 | 2162.0 ± 1332.9 | 9.0 – 6711.0 |
| Cholesterol | ml dl<sup>-1</sup> | 32 | 32 | 0 | 219.8 ± 64.6 | 18.5 – 340.7 |
| Phospholipids | ml dl<sup>-1</sup> | 32 | 32 | 0 | 347.4 ± 106.4 | 41.9 – 526.4 |
| Non-esterified fatty acids | ml dl<sup>-1</sup> | 30 | 30 | 0 | 0.7 ± 0.2 | 0.3 – 1.3 |
| PC song + PC condition | NA | 41 | 41 | 0 | NA |  |
| PC song + ddRAD | NA | 101 | 81 | 20 | NA |  |
| PC condition+ ddRAD | NA | 49 | 49 | 0 | NA |  |
| PC song + PC condition + ddRAD | NA | 39 | 39 | 0 | NA |  |

**Figure 2:**
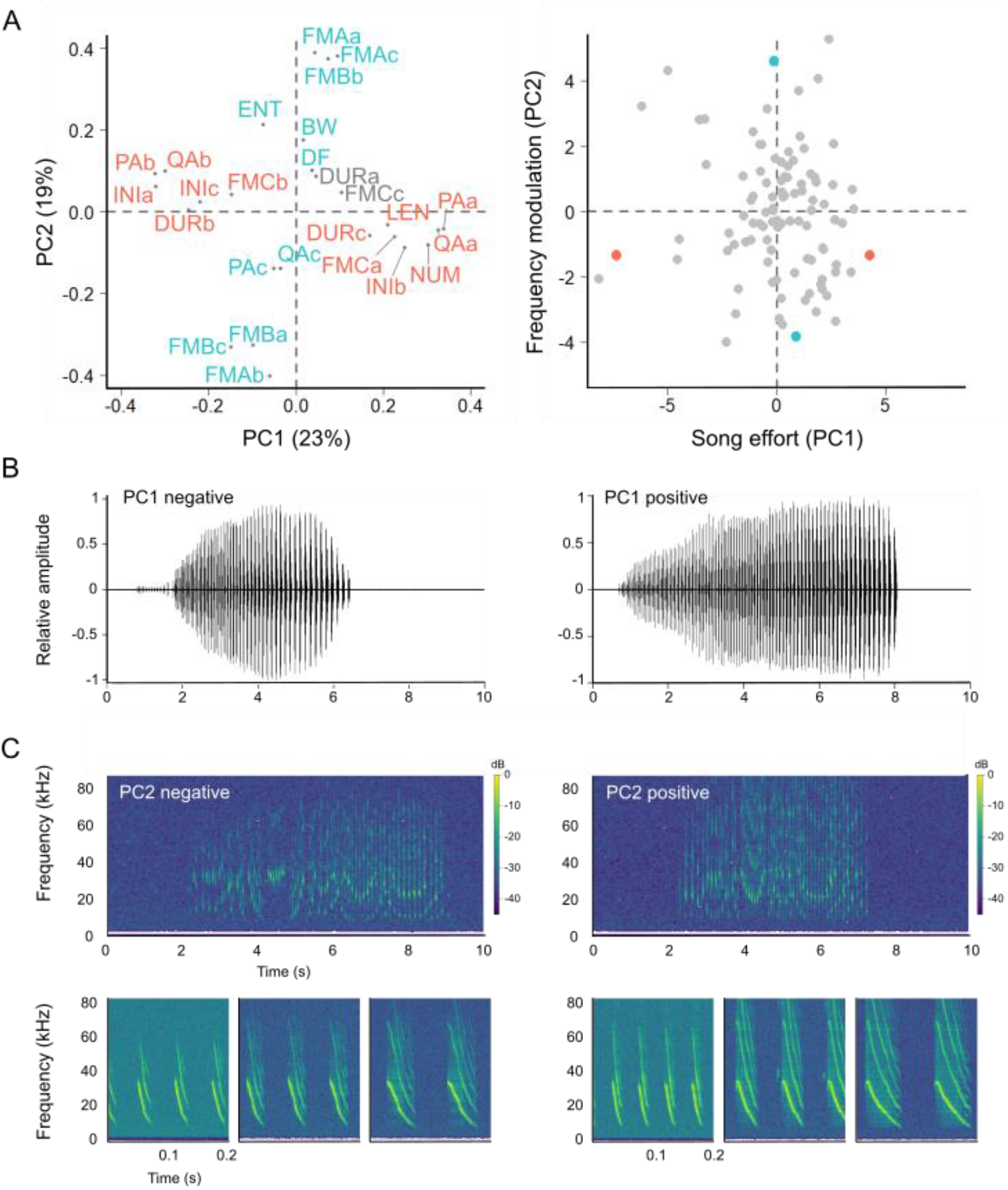
Acoustic variation in songs of wild singing mice. *(A)* Biplots from PCA of *Scotinomys* song. *Left*, biplot showing how variables loaded in PC space. In red, parameters loading strongly on PC1 (“song effort”). In blue, parameters loading strongly on PC2 (“spectral characteristics”). *Right*, individual singers plotted in PC space. Points colored red and blue are individuals that produced the songs plotted in B and C, respectively. *(B)* Variation in song effort. *Left*, song loading strongly and negatively on PC1. *Right*, song loading strongly and positively on PC1. *(C)* Variation in spectral characteristics. Top panels depict full spectrograms of songs with lower panels depicting 0.1 ms of song taken at 1 s, 3 s, and 5 s. *Left*, song loading strongly and negatively on PC2. *Right*, song loading strongly and positively on PC2.

## Results

### Phenotypic sampling

We sampled a population of singing mice from San Gerardo de Dota, Costa Rica (Fig. 3, Table 1), taking a variety of measurements of song and condition from these wild-caught animals. Summary statistics, sample sizes, and breakdown by sex for all measured phenotypes are reported in Table 2.

**Figure 3:**
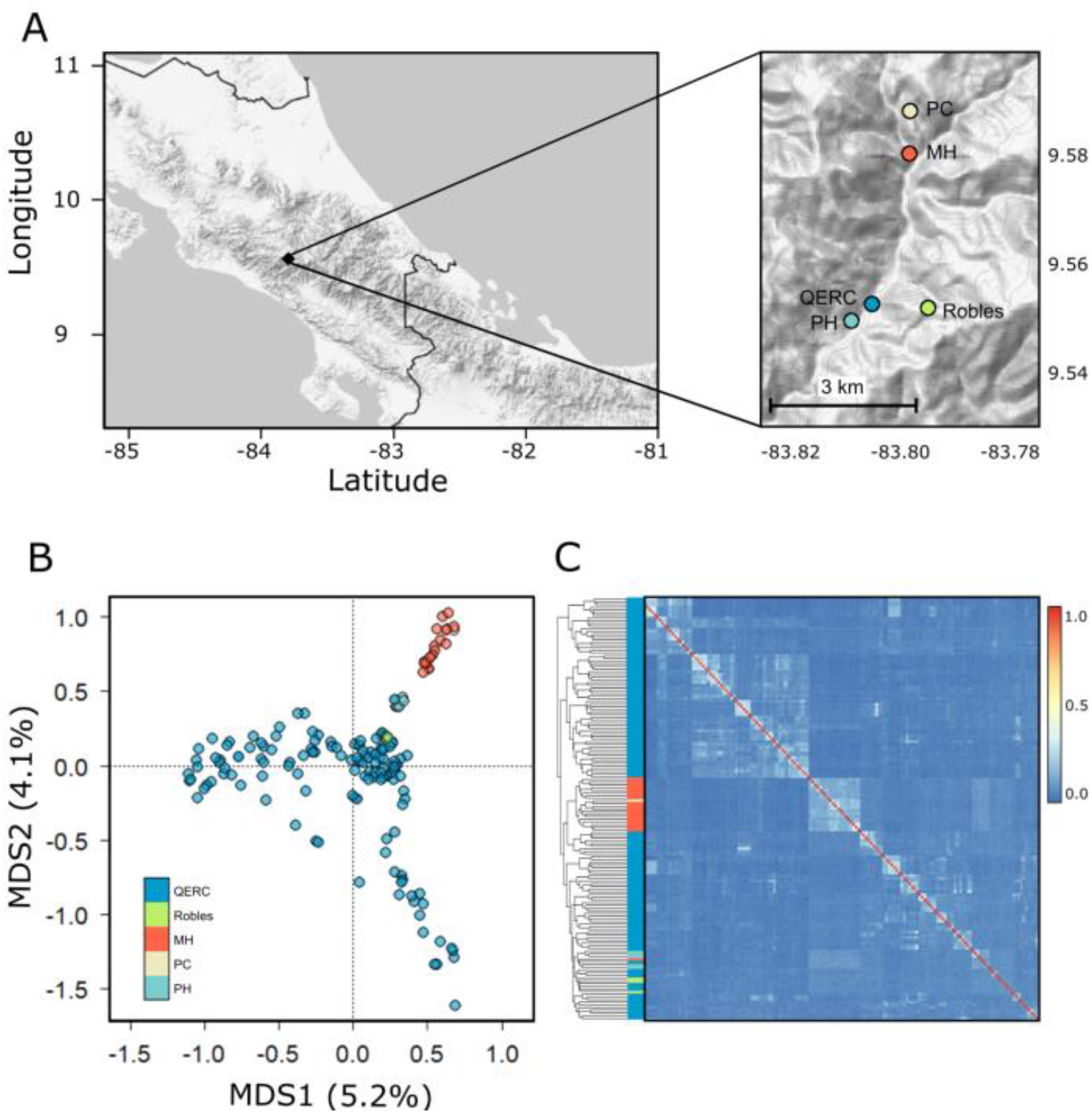
Geographic structure of relatedness in a wild population of singing mice. *(A)* Map of Costa Rica with zoomed-in panel of trapping sites within the valley of San Gerardo de Dota. *(B-C)* Colored by site from *A*. *(B)* Biplot of first two PCs from PCoA of genetic distance between individual mice. *(C)* Heatmap of genetic relatedness matrix (GRM) for 157 mice. Hierarchical cluster branches annotated by trapping site as in *A*. See Table 1 for elevation data and sample sizes.

We successfully recorded long songs and spontaneous song rate (*i.e.* number of songs in two hours) from 106 and 129 mice, respectively (Table 2). Mice with at least one song were included in principal components analysis (PCA) of acoustic variation, which revealed two latent variables, PC1 (“song effort”, 23.1% of overall variance), which explained individual variation in amplitude and duration, and PC2 (“spectral characteristics”, 19.1%), which explained individual variation in frequency characteristics of song, particularly note shape (Fig. 2, Table S1).

We regressed body mass (RBM) as a common measure of body condition in 165 mice and measured anogenital distance (AGD) in 159 mice. We took plasma samples from each of the 165 mice, but only male plasma was processed; of these, 78 samples had enough plasma to complete at least one assay. All plasma measures fell within the range of values reported for laboratory mice except adiponectin, an adiposity hormone involved in glucose and fatty acid metabolism (*Mus*: 3000–15 000 ng ml^−1^, Friedman and Halaas, 1998; *Scotinomys*: 50–350 ng ml^−1^, Burkhard et al., 2018). Fifty-two samples met the requirements to be included in PCA of body condition (see Methods), which revealed three latent variables. PC1 (“PC1 condition”, 34.5% of total variation) described individual variation in glucose, fatty acids, and insulin, which we interpreted as metabolic responses to recent feeding. PC2 (“PC2 condition”, 22.2%) described variation in residual body mass (RBM) and adiponectin, an adiposity hormone that modulates glucose and lipid metabolism; while PC3 (“PC3 condition”, 17.2%) described variation in leptin, an adiposity hormone that helps regulate body weight, and adiponectin (Table S2). We interpreted these latter two composite variables as putatively stable indicators of long-term body composition and regulation of energy stores.

### Variant calling

We used ddRAD-sequencing (Peterson et al., 2012) to genotype 162 mice. Five samples had poor coverage and were excluded from downstream analyses. We demultiplexed and trimmed reads using the *process_radtags* command from the Stacks pipeline (v1.46.0; Rochette and Catchen, 2017), then mapped trimmed reads to the *S. teguina* reference genome (annotated, unpublished) using Bowtie 2 (v2.3.2; Langmead and Salzberg, 2012). After pruning the remaining 157 genotypes for minor allele frequency (MAF) > 0.1 and for sex-linked markers using VCFTOOLS (v0.1.15; Danecek et al., 2011) and ANGSD (v0.929-13; Korneliussen et al., 2014), we retained 31,003 SNPs.

### Population structure and individual relatedness

Because population structure can influence estimates of heritability, we first examined population genetic structure (Fig. 3). Permutational multivariate analysis of variance (PERMANOVA) revealed low but statistically significant genetic differentiation between trapping sites (*R*^*2*^ = 0.027, *P* = 0.001), and principal coordinate analysis of the genotypic data revealed distinct but overlapping genetic clusters explained by trapping site (Fig. 3b). Consistent with this interpretation, we estimated generally low *F*ST between sites (range: 5.7E-06 – 0.13) and found *F*ST increased with geographic, though not elevational, distance (two-sided Mantel test; geographic distance: *r* = 0.68, *P* = 0.02; elevational distance: *r* = −0.05, *P* = 0.91; Table 3). We next estimated relatedness of individuals within our sample. Most pairwise genetic relationships were the equivalent of unrelated individuals, but we also uncovered stronger genetic relationships equivalent to half-sibling and full-sibling relatedness (Fig. 3c).

**Table 3:**
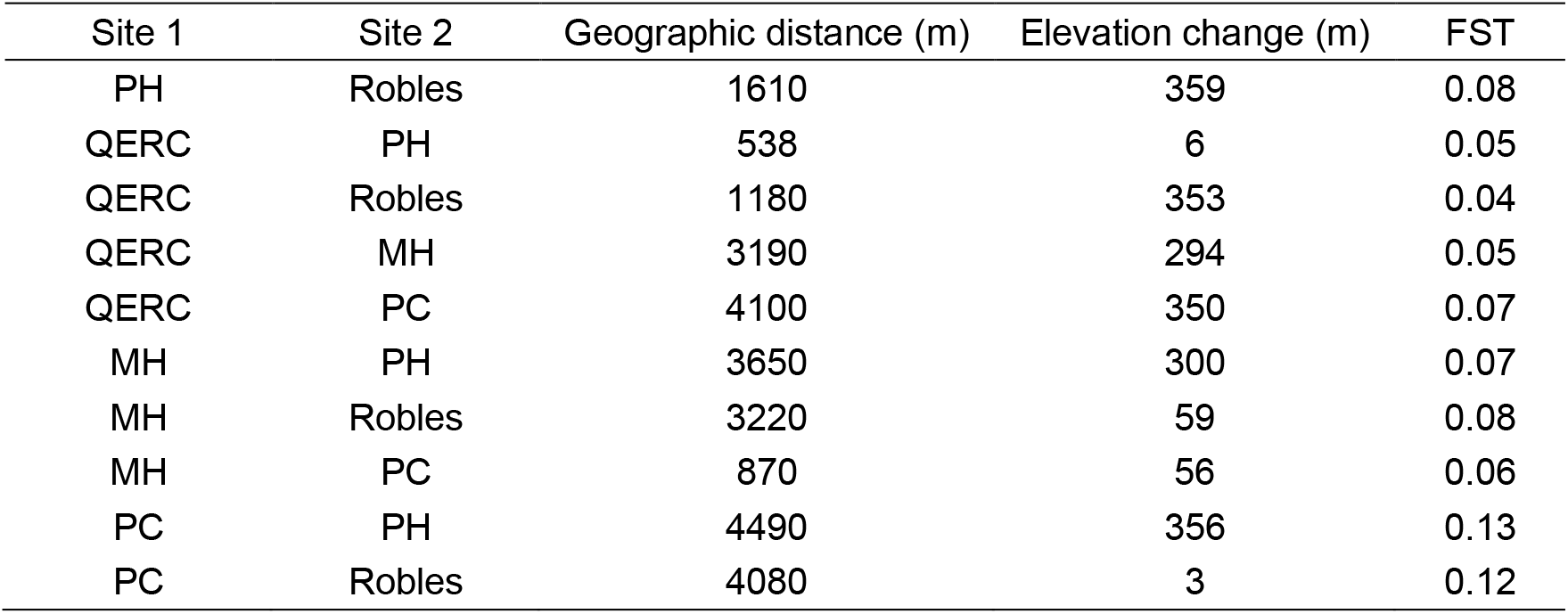
Trapping site statistics. Geographic distance and elevational change calculated from GPS waypoints.

### Repeatability of acoustic variation

We estimated repeatability of acoustic characteristics by calculating intraclass correlation coefficients (ICC). Repeatability ranged from negligible (ICC < 0.2) to high (ICC ≥ 0.5; Table S3) and was generally greater for spectral song descriptors than for those related to song effort (spectral: mean *ICC* = 0.36, range = 0 – 0.76; effort: *ICC =* 0.21, 0 – 0.45). This pattern became more pronounced when considering only whole-song descriptors (*i.e.* song length, dominant frequency; Table S3), with whole-song spectral measures having greater mean repeatability than whole-song measures of effort (spectral: mean *ICC* = 0.32, range = 0.43 – 0.76; effort: *ICC* = 0.32, 0.30 – 0.33; Table S3, Fig. 4). The top three most repeatable aspects of song (*ICC* ≥ 0.6) were spectral characteristics: song mean bandwidth (*ICC* ± SE; BW: 0.76 ± 0.02), minimum frequency (minHz: 0.76 ± 0.03), and the starting frequency of the first note of a song (FMc_c: 0.61 ± 0.04). Entropy, a whole-song description of tonality (0.59 ± 0.04), maximum frequency (maxHz: 0.56 ± 0.05), and FMb_c, starting note slope (0.52 ± 0.04) were the other measures with ICC ≥ 0.5. Twelve parameters, including dominant frequency and song length, were modestly repeatable (0.5 > *ICC ≥* 0.2), while the remaining 13 parameters had negligible repeatability.

**Figure 4:**
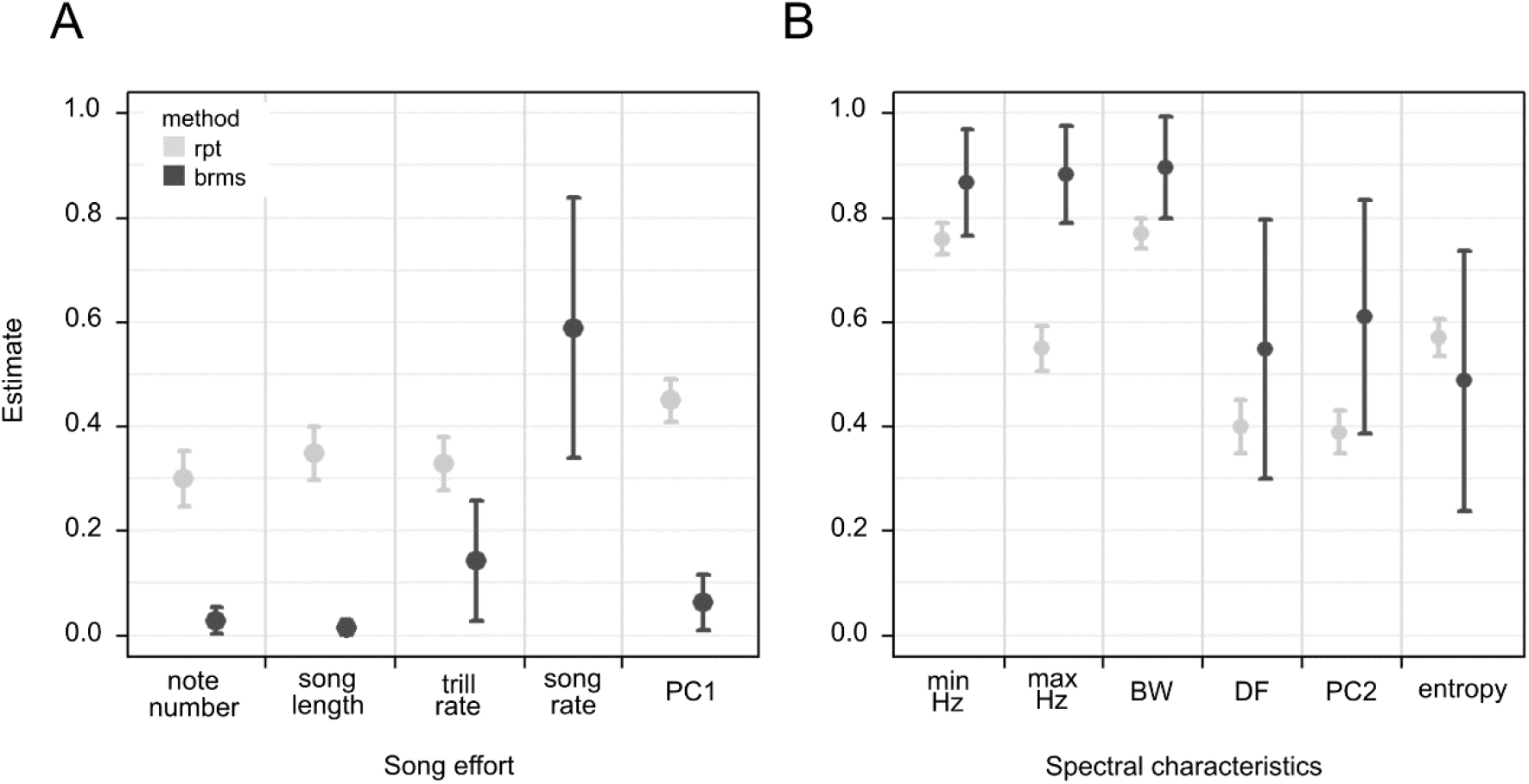
Heritable and repeatable variation in songs. *(A)* Repeatability (rpt) and heritability (brms) estimates from different models for song effort characteristics. *(B)* Estimates for spectral characteristics. Error bars indicate standard errors.

### Heritability of song and condition

We estimated heritability by fitting animal models using maximum likelihood (ML) estimation implemented using standalone software *GCTA* (Yang et al., 2011) and using Bayesian Markov chain Monte Carlo methods implemented with R-package *brms* (Bürkner, 2018, 2017). Heritability estimates derived from *GCTA* and *brms* were generally consistent for the same phenotypes with a notable exception, PC1 condition (*GCTA*: *h*^*2*^ = 0.96 ± 0.59, *brms*: *h*^*2*^ = 0.07 ± 0.07; Table 4, Table S4). Such disagreements result from how the two approaches deal with sparse data—for example, due to low levels of true heritability or small sample size. In these cases, ML estimation tends to be extremely variable and may report values that are either very large or zero (*e.g.* Beerli, 2006), while Bayesian approaches, given appropriate priors, offer more reliable results, as demonstrated by analyses of both empirical and simulated data (Beerli, 2006; Charmantier et al., 2011; Van de Schoot et al., 2015; Villemereuil et al., 2013). Credible intervals from Bayesian analysis are easily interpretable and provide distributional information indicating the most probable value and associated certainty (de Villemereuil, 2019; Kruschke and Liddell, 2018). For these reasons, we report here only heritability estimates (± SE) and 95% highest posterior density credible intervals (HPDI) from Bayesian models. For comparison, GCTA results are reported in Supplemental Files (Table S4).

**Table 4:**
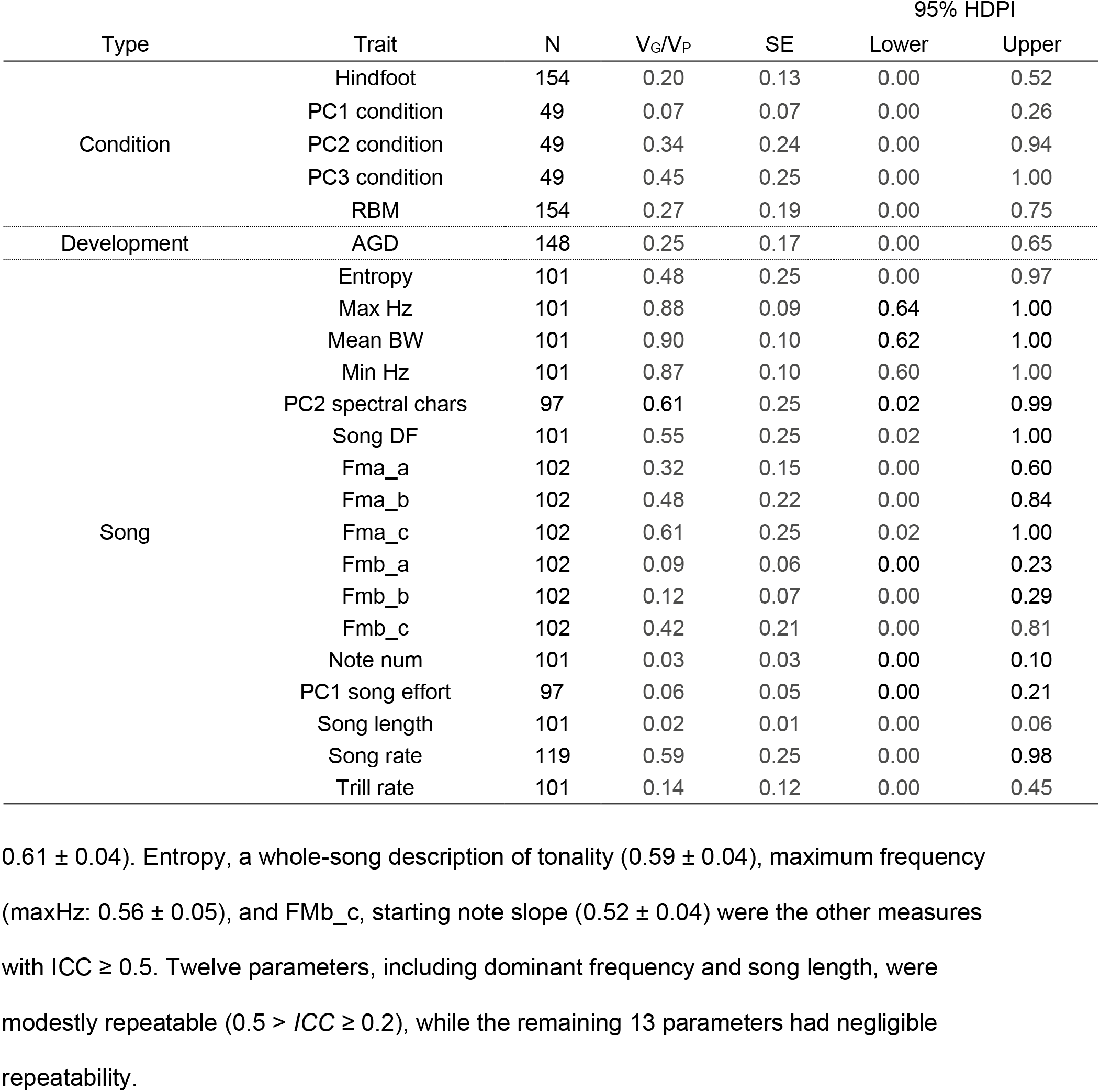
Heritability estimates for song and condition phenotypes. 95% HPDI = 95% highest posterior density credible interval.

#### Patterns of heritable variation in songs

Heritability estimates for whole-song measures related to song effort (*h*^*2*^ range: 0.02 – 0.59, mean: 0.19) were generally lower than estimates for whole-song spectral measures (*h*^*2*^ range: 0.48 – 0.90, mean: 0.57; Fig. 4, Table 4). PC composite variables followed this pattern, with PC1 having negligible heritability (*h*^*2*^ = 0.06 ± 0.05) and PC2 having high heritability (*h*^*2*^ = 0.61 ± 0.25). Mean song bandwidth, maximum song frequency, and minimum song frequency were the most heritable acoustic traits, with *h*^*2*^ ≥ 0.8, while song length and note number were the least heritable traits, with *h*^*2*^ ≤ 0.05. One notable exception to the general pattern was song rate, the number of songs produced in two hours, which had high heritability (*h*^*2*^ = 0.59 ± 0.25).

We also examined the repeatability and heritability of several measures describing note shape (*i.e.* note curvature, FMa_X; note slope, FMb_x) and how shape changes over the course of a song (Fig 5); for example, FMa_A describes how rapidly note-to-note differences in note curvature happen over course of song, FMa_B describes the change in note curvature between first and second notes, and FMa_C describes the note curvature in first note in song (Campbell et al., 2010). We estimated intermediate to high repeatability and heritability for FMx_C terms; by contrast, we found low to intermediate repeatability and heritability for FMx_B terms and negligible to low repeatability and heritability for FMx_A terms (Tables 4, S3).

**Figure 5:**
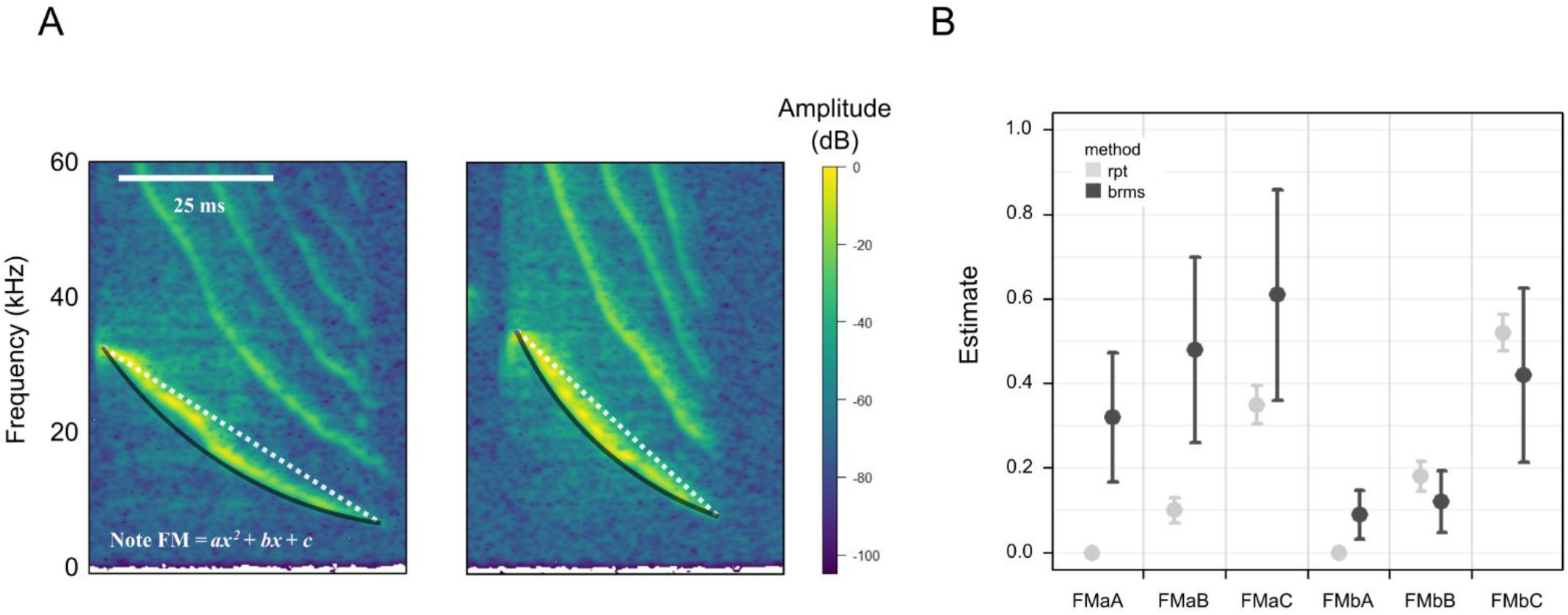
Heritability of note shape. *(A)* Close-ups of a single note over 50 ms. *Left*, note with low FMb and FMa (shallow note slope and curvature). *Right*, note with high FMb and FMa (steep note slope and curvature). Black lines trace note curvature; dotted white lines indicate note slope. *(B)* Repeatability (rpt) and heritability (brms) estimates from different models for song effort characteristics. Error bars indicate standard errors.

#### Patterns of heritable variation in body condition and morphology

Condition and morphological measures ranged in heritability from low to intermediate heritability (*h*^*2*^ range: 0.07 – 0.45, Fig. 6a, Table 4). Anogenital distance (AGD), a morphological indicator of developmental exposure to androgens, had moderate heritability (*h*^*2*^ = 0.25 ± 0.17). Hindfoot length and RBM also had moderate heritability (hindfoot: *h*^*2*^ = 0.20 ± 0.13; RBM: *h*^*2*^ = 0.27 ± 0.19). Measures of long-term body composition (*i.e.* PC2 and PC3 condition) were more heritable (PC2 condition: *h*^*2*^ = 0.34 ± 0.24; PC3 condition: *h*^*2*^ = 0.45 ± 0.25). Finally, short-term fluctuations in nutrients (PC1 condition) had low heritability (*h*^*2*^ = 0.07 ± 0.07).

**Figure 6:**
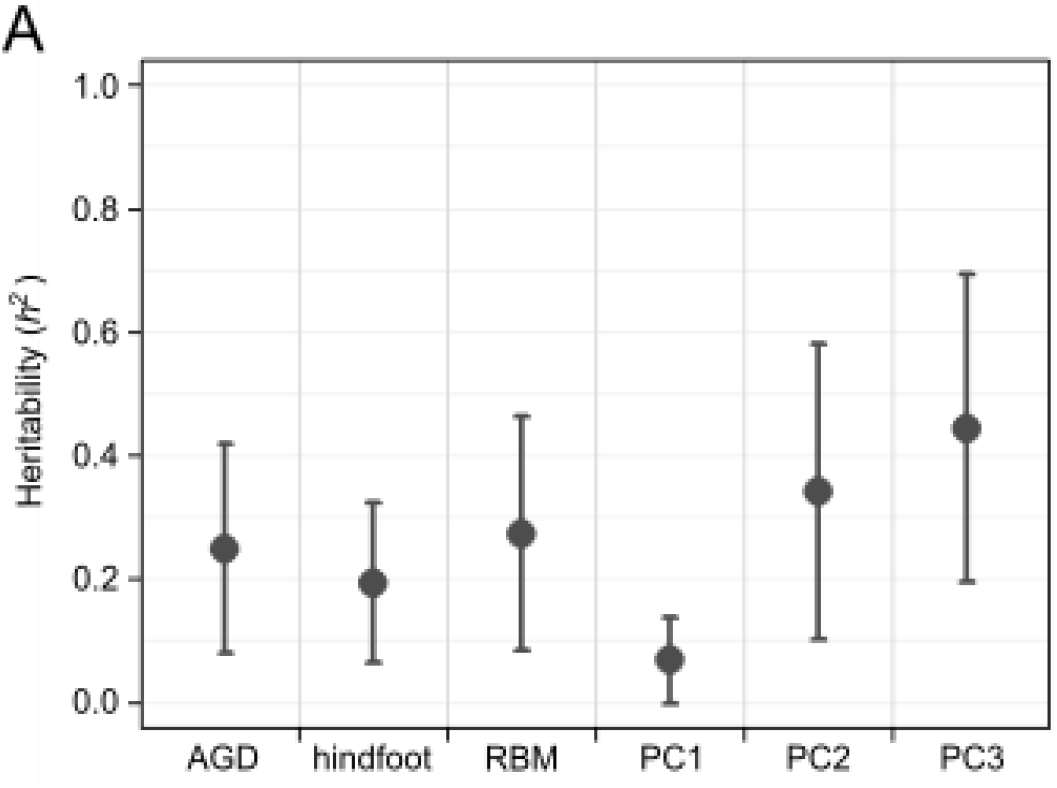
Heritability of condition. Error bars indicate standard errors.

### Correlations between body condition and singing behavior

#### Phenotypic correlations of condition and song

We next examined phenotypic correlations of condition and song by fitting mixed models. Forty-one male mice had both song and PC condition data and could be included in linear or generalized linear mixed models for condition-dependence (Tables 2, S5). Acoustic characteristics related to song effort were predicted by both body size and adiposity measures of condition: males with high PC2 condition scores sang longer songs with more notes than individuals with low PC2 condition scores (song length: *β*_*PC2con*_ = 0.32, *SE* = 0.10, *P* = 0.002; note number: *β*_*PC2con*_ = 0.32, *SE* = 0.10, *P* = 0.001, Fig. 6b). Similarly, males with high PC3 condition scores (high leptin, low adiponectin) produced more songs in the span of two hours than males with low PC3 condition scores (song rate: *β*_*PC3con*_ = 0.21, *SE* = 0.10, *P* = 0.04, Fig. 6c). Interestingly, we did not find a statistically clear effect of body size or adiposity measures on PC1 song effort, but we did find a clear relationship between song effort and AGD, an indicator of phenotypic masculinity (song effort: *β*_*agd*_ = 0.76, *SE* = 0.36, *P* = 0.04).

While increases in PC3 condition scores also predicted modest increases in song maximum frequency (max Hz: *β*_*PC3con*_ = 0.07, *SE* = 0.13, *P* = 0.007) and modest decreases in song minimum frequency (min Hz: *β*_*PC3con*_ = −0.04, *SE* = 0.01, *P* < 0.001), other spectral characteristics were not strongly predicted by any other measure of body condition (Table S5). Finally, PC1 condition, variation in immediate metabolic responses to recent feeding, did not predict any measure of song (Table S5).

#### Co-heritability of condition and song

We estimated genetic correlations for combinations of song effort characteristics (song length, note number, song rate) and condition measures (PC2, PC3) that were substantially phenotypically correlated (Table S5). We also investigated genetic correlations between song effort characteristics and RBM; this comparison allowed us to examine the utility of the traditional body condition metric and afforded us greater statistical power. Correlations ranged from −0.19 to 0.61 and were generally larger in magnitude for traits that were more strongly phenotypically correlated (Tables 5, S5). Total songs and PC3 of condition had the largest genetic correlation (*r* = 0.61 ± .29); this was unsurprising given the intermediate to high heritability estimates for each trait. In all cases, however, the 95% HPDI included the region of practical equivalence (ROPE) around 0 (Kruschke and Liddell, 2018); thus, we did not find clear statistical evidence for significant genetic correlations in our study.

**Table 5:** Genetic correlations (*r*_*G*_) between condition and song. 95% HPDI = highest posterior density credible intervals.

| Trait 1 | Trait 2 | N | $r_G$ | SE | 95% HPDI | |
| --- | --- | --- | --- | --- | --- | --- |
|  |  |  |  |  | lower | upper |
| call length | RBM | 80 | 0.14 | 0.15 | -0.17 | 0.43 |
| note number | RBM | 80 | -0.01 | 0.34 | -0.65 | 0.66 |
| total songs | RBM | 118 | 0.19 | 0.35 | -0.7 | 0.87 |
| call length | PC2 condition | 31 | 0.35 | 0.2 | -0.07 | 0.69 |
| note number | PC2 condition | 31 | 0.02 | 0.34 | -0.63 | 0.64 |
| total songs | PC3 condition | 49 | 0.61 | 0.29 | -0.12 | 0.98 |

## Discussion

In this study, we used genome-wide data to investigate heritable variation in a variety of phenotypes related to advertisement display and body condition in a Costa Rican population of Alston’s singing mouse (*Scotinomys teguina*). We obtained 31,000 SNPs from ddRAD-seq that we then used to estimate genome-wide relatedness, finding genetic relationships ranging from unrelated (rel ≅ 0) to the equivalent of full siblings (rel ≅ 0.5). We next calculated SNP-based heritability for various characteristics of singing behavior and measures of body condition, confirming results with both frequentist and Bayesian modeling. We found that measures of repeatability, a common surrogate for heritability (Boake, 1989), accurately predicted patterns of heritability: characteristics related to song effort, which were modestly repeatable, had low to moderate heritability, whereas spectral characteristics were often highly repeatable and heritable. We estimated intermediate heritability for patterns of body adiposity, while the heritability of insulin and circulating nutrients was negligible. Adiposity and song effort exhibited a strong phenotypic correlation, which was accompanied by a surprisingly high estimate of their co-heritability (*r* = 0.61). We now explore these findings in more depth.

### Heritability and repeatability of singing behavior

*Scotinomys* songs vary in a wide range of acoustic attributes, and a principal components analysis of acoustic features suggested two major dimensions of song variation: those related to duration and amplitude, which we consider “song effort”; and those describing spectral characteristics, specifically, the frequency modulation that occurs over the course of a song (Fig 2, 4). Although heritability varied greatly from feature to feature, ranging from values near 0.0 to above 0.8, there were systematic patterns to the heritability of songs and their attributes (Table 5).

Among our various measures, whole-song spectral variation (dominant, maximum, and minimum frequency; mean bandwidth, entropy) was both highly repeatable and heritable. We hypothesize that this heritability reflects anatomical variation or low-level motor control. For example, dominant frequency (*h*^*2*^ = 0.55 ± 0.25; *ICC* = 0.43 ± 0.05) is a property of both the vocal tract and adjustments to its length or shape (Fitch, 2006; *e.g.* via lip contractions in humans, Story et al., 2001; tongue placement or bill gape in songbirds, Langin et al., 2017, Nelson et al., 2005; laryngeal descent in humans and deer, Fitch and Reby, 2001). Maximum frequency (*h*^*2*^ = 0.88 ± 0.1; *ICC* = 0.56) is likely a function of the cricothyroid, a muscle controlling the length and tension of vocal folds (Jürgens, 2002; Riede and Brown, 2013). In muroid mice that produce laryngeal whistles, variation in structures like the vocal folds and the ventral pouch are proposed to shape interspecific variation in dominant frequency (Riede et al., 2017; Riede and Pasch, 2020; Smith et al., 2020). At a phylogenetic scale, spectral properties are often constrained by body size, a relationship generally mediated by scaling of vocal structures (Gillooly and Ophir, 2010; Ophir et al., 2010; Ryan and Brenowitz, 1985). The relationship between body size and frequency is often observed within species as well, such as in red deer (Reby and McComb, 2003), anurans (Gerhardt, 1982), songbirds (Ryan and Brenowitz, 1985), loons (Mager et al., 2007), and humans (Pisanski et al., 2014). It is by no means universal, however (*e.g.* Martin et al., 2017), and we did not observe a relationship in our sample.

While whole-song spectral characteristics were highly heritable, we also found substantial heritability in frequency modulation. Six related variables strongly loaded on PC2; together, these variables describe the frequency modulation that characterizes a note and how it changes over the course of a song (Figs. 2, 5). To appreciate how these are related, imagine a single note composed of a downward frequency sweep that begins and ends at characteristic frequencies defined by the performance of the vocal tract. A given note may proceed linearly from the beginning frequency to the end frequency with a relatively shallow slope, or may change rapidly in frequency initially, and then flatten out (Fig. 5a). These two parameters are the starting slope and curvature of a note (*i.e.* FMb, and FMa, the linear and quadratic terms in the equation: *Frequency* = *FMc* + *FMb* × *time* + *FMa* × *time*^2^; Figs. 1, 5). We find that the slope and curvature of notes are strongly correlated with one another, and change systematically over the course of the song. Starting note shape (FMb_C, FMa_C) exhibits high repeatability and heritability (Tables 4, S3). Unlike many other spectral features of the song, note shape reflects the temporal dynamics of muscle activity over the course of a note. Such timing must reside in central pattern generators (CPGs), such as those vocal, orofacial and respiratory CPGs thought to reside in the brainstem (Zheng, 2020). In singing mice, laryngeal and jaw muscles important for vocalization seem to be coordinated by a putative CPG within the reticular formation (Zheng et al., *manuscript*).

The fact that note shapes vary by individual and are highly heritable is also consistent with prior suggestions that they contribute to identity signaling in this species (Burkhard et al., 2018). Other studies have found that ecological contexts can favor individually distinctive social signals (Burt and Beecher, 2008; Sheehan and Tibbetts, 2010; Stoddard and Beecher, 1983). If this were true, it would predict that not only could animals use note shape and other spectral features to recognize individuals, but that balancing selection may actively promote diversity of neural activity patterns within CPGs. These results highlight a need to examine whether note shapes are in fact used in individual recognition.

In contrast to spectral features, we found low heritability for most song-effort traits. Our PC1 of song variation, which we labelled song effort, was low both in its repeatability and heritability (ICC = 0.21, *h*^*2*^ = 0.06). Measures like song length, note number, and trill rate (*ICC* < 0.35) all had negligible heritability (*h*^*2*^ < 0.1). One explanation for this pattern is that highly plastic behaviors, which are shaped by social context and experience, may be less heritable than traits constrained by morphology (Boake, 1989; Mousseau and Roff, 1987). This phenomenon has been documented in other natural populations (*e.g.* three-spined stickleback: Dingemanse et al., 2009; island-adapted deer mice: Baier et al., 2019) and confirmed by meta-analyses of behavioral and morphological datasets (Mousseau and Roff, 1987; Stirling et al., 2002; but see Jensen et al., 2003; Dochtermann et al., 2019). This interpretation is consistent with evidence from singing mice suggesting that song effort is highly sensitive to social context and motivation. Male singing mice increase song rate in response to winning fights (George, 2014), encounters with females (Fernández-Vargas et al., 2011; Hooper and Carleton, 1976), and simply hearing another animal vocalize (Hooper and Carleton, 1976; Pasch et al., 2013). Singing mice also exhibit dynamic “turn-taking” counter-singing behavior, rapidly modifying the starts and stops of songs during social interactions (Okobi et al., 2019). Similar relationships between vocal effort and social experience are well documented in other taxa, particularly in avian (Catchpole and Slater, 2008; Podos and Cohn-Haft, 2019) and anuran species (Bernal et al., 2009; Gerhardt, 1991; Ryan, 1985), in wild muroid rodents (Fernández-Vargas, 2018 a; Fernández-Vargas, 2018 b), and in other mammals (Briefer, 2012; Gouzoules and Gouzoules, 2000), including humans (Bachorowski, 1999). Male music frogs dynamically adjust calling efforts based on perceived sexual attractiveness of rivals (Fang et al., 2014), comparable to male responses to competition in gray treefrogs (Gerhardt, 1991) and túngara frogs (Bernal et al., 2009; Ryan, 1985). Likewise, pair-bonded California mice increase their use of aggressive “bark” vocalizations towards recently unfaithful partners (Pultorak et al., 2018), a behavioral response with uncanny parallels to non-verbal human expressions of anger (Sauter et al., 2010).

Taken together, our data suggest that as we move from the peripheral vocal organs, through low-level motor control, and finally to the higher-level neural centers that govern motivational state and dynamic social interactions, aspects of song become progressively less heritable. This seems to be a major finding of our analysis, but there are two anomalous aspects of our data that merit some further discussion. The first is that stable differences in song duration, trill rate, and related characteristics that seem negligibly heritable in our sample are nevertheless distinct among populations of singing mice (Campbell et al., 2010; Hooper and Carleton, 1976); these differences persist in laboratory-reared mice (Campbell et al., 2014), suggesting a heritable basis for lineage differences. One possibility is that meaningful heritability is present in these measures, but our power is insufficient to detect it. A second possibility is that low heritability in contemporary populations may be a by-product of past selection. Local adaptation, for example, would preserve between-lineage heritability while reducing within-population genetic variance (Falconer, 1981; Mousseau and Roff, 1987). Future work will investigate the relationship between acoustic and genetic variation across populations of singing mice. The second anomaly in our findings is that song rate, which we interpret as a motivational aspect of song, is surprisingly heritable (*h*^*2*^ = 0.59 ± 0.25). We recently reported that manipulating perceived body condition by injecting leptin increases song rate, but not song duration. Perhaps song rate is particularly strongly influenced by trade-offs associated with body condition – a hypothesis that is supported by phenotypic correlations between the two dimensions (discussed below).

### Heritability of condition

Body condition holds evolutionary significance for two major reasons: it is thought to be tightly linked to fitness, and it is integral to many models of signal evolution. As a result, its heritability is also of evolutionary interest. While fitness-related traits are generally expected to have low heritability (Falconer, 1981; Mousseau and Roff, 1987), it is not immediately obvious whether condition should follow this pattern. First, certain indices of condition may not be as predictive of evolutionary fitness as others, or may predict different aspects of fitness; for instance, in damselflies, body weight is negatively correlated with parasite-mediated mortality but has no relationship with reproductive success (Rolff et al., 2002). Further, condition may be variably affected by contributions of environmental and genetic variation underlying its many physiological and morphological constituents. Lastly, if condition is highly polygenic due to its dependence on a large number of other aspects of phenotype, the high “mutational target size” of the trait could allow heritable variation to persist in the face of selection – an argument that has been used to suggest that elaborate displays can serve as indicators of underlying genetic viability (Rowe and Houle, 1996).

In the present study, we described body condition with measures of residual body mass and various circulating nutrients and metabolic hormones, using PCA to define major dimensions of variation. PC1 described variation in glucose, fatty acids, and insulin, which we interpreted as immediate metabolic responses to recent feeding or fasting. PC2 and PC3 each reflected patterns of adiposity: PC2 described variation in body size and adiponectin, an adiposity hormone involved in glucose and fat digestion, while PC3 described variation in adiponectin and leptin, an adiposity hormone regulating body weight. Thus, we interpreted both PC2 and PC3 of condition as putatively stable indicators of body composition.

We found low heritability for circulating nutrients and insulin (PC1 condition: *h*^*2*^ = 0.07), a result that is consistent with these measures as dynamic responses to recent feeding. Relative to the moderate heritability of RBM, stable indicators of body composition (PC2 and PC3 condition) were higher in heritability (Table 4, Fig. 6) but were lower than expected. Evidence from humans and laboratory rodents suggests a high heritability of adiposity- and metabolism-related traits, including body fat distribution (Schleinitz et al., 2014), percent body fat (Wuschke et al., 2007), fat type (Ferrannini et al., 2016), and metabolic rate (Pettersen et al., 2018); indeed, twin studies report higher heritability of human body fat (*h*^*2*^ = 0.59-0.63, Schousboe et al., 2004; *h*^*2*^ = 0.71-0.75, Lehtovirta et al., 2010) and body mass index (BMI; *h*^*2*^ = 0.58-0.63, Schousboe et al., 2004; *h*^*2*^ = 0.63-0.85, Allison et al., 1996) than values we report here. However, an important distinction between human or lab rodent work and this study is that the former significantly reduces environmental variance by design whereas our focal population has been subject to natural environmental conditions.

### Condition dependence

Elaborate displays can be costly to express, and are thought to rely on body condition (Bradbury and Vehrencamp, 1998; Cotton et al., 2004; Johnstone, 1993). Condition dependence thus plays an important role in individual signaling decisions, determining the expression and intensity of labile signals, in addition to serving as a key mechanism for many models of signal evolution (Grafen, 1990; Kirkpatrick and Ryan, 1991; Pomiankowski, 1987; Zahavi, 1975).

Condition-dependence could also lead to genetic covariance between loci affecting condition and display, a critical prediction of good-genes models of signal evolution (Kirkpatrick and Ryan, 1991; Rowe and Houle, 1996; Tomkins et al., 2004).

We tested whether major dimensions of song were phenotypically corelated with measures of condition. As expected, stable spectral characteristics, putatively constrained by variation in vocal structures and morphology, lacked strong correlations with measures of body condition. Conversely, we found significant relationships between elements of song effort, including song length, vocalization rate, and note number, and stable measures of body composition (PC2 and PC3 of condition). These findings are consistent with previously reported relationships between body condition and song in singing mice (Burkhard et al., 2018; Pasch et al., 2011b) and with general patterns described in other taxa (*e.g.* birds: Houtman, 1992, Lampe et al., 1994; crickets: Thomson et al., 2014). For example, Thomas and Cuthill (2002) demonstrated that European robins (*Erithacus rubecula*) song rate was positively correlated with body mass, such that increases in body mass (*i.e.* increased fat reserves) predict more frequent singing behavior (Thomas and Cuthill, 2002). Likewise, greater body mass predicts more elaborate repertoires and better performance in Java sparrows (*Lonchura oryzibora*) (Kagawa and Soma, 2013). Possible explanations for this general phenomenon include the hypothesis that conspicuous song, along with other labile displays, may be energetically costly to produce (Barske et al., 2013; Ward et al., 2003) or may elicit the energetic costs of predator evasion (Zuk and Kolluru, 1998). Signaling decisions are thus likely modulated not only by social context but by self-assessment of body condition (Santori et al., 2020), perhaps through interoceptive cues, like adiponectin or leptin, that signal status of energetic reserves (Havel, 2001).

Both song rate and PC3 of condition (the adiposity hormones leptin and adiponectin) were remarkably heritable (Table 5), and our estimate for the genetic correlation between the two suggests that roughly 60% of the heritability in song rate could be predicted by this measure of condition (Table 6). This estimate is surprisingly large – it is comparable, for example, to published genetic correlations between closely related psychiatric conditions like schizophrenia and bipolar disorder (r_g_ = 0.64, Lee et al., 2013). An important caveat, however, is that the credible intervals overlap the region of practical equivalence around 0 (Kruschke and Liddell, 2018), indicating that we lack the power to precisely estimate the genetic correlation. Our findings indicate, however, that it would be worthwhile to generate a more precise estimate of this genetic correlation. Such studies should focus on hormonal indicators of condition, and should include a substantially larger number of animals. What constitutes a reasonable sampling effort will vary by study system, its demography, and its natural history (eg Bérénos et al., 2014; Villemereuil et al., 2013).

Finding a genetic correlation is necessary for the good genes hypothesis, but it is not sufficient. The idea that an expensive trait could be an index for heritable variation in condition is plausible, but we would expect genetic correlations to be present even if female choice plays no role in the evolution of effort—because males must make these decisions in order to balance risks and rewards in general. Our approach, however, provides a way to generate meaningful measures of body condition in the wild, as well as meaningful measures of heritability and co-heritability. Such data will be essential to disentangling the role of direct benefits – such as males maximizing the outcomes of trade-offs, and females avoiding parasites – and indirect benefits, such as minimizing the genetic burden of deleterious mutations passed on to offspring.

## Conclusions

For decades, the challenges of studying heritability in non-model and wild species has compelled many behavioral ecologists and field biologists to either strategically ignore the genetic mechanisms underlying traits (the “phenotypic gambit”; Grafen, 1984; Rittschof and Robinson, 2014) or focus instead on more tractable but less informative alternative measures, like repeatability (Boake, 1989). Despite the obvious logistical advantages of lab estimates of heritability, field estimates are more ecologically meaningful. Lab conditions can reduce genetic variation and environmental variation, which may lead to under- or overestimation of heritability (Orengo and Prevosti, 1999; Roff and Simons, 1997; Schneider et al., 2011; Simons and Roff, 1994; Stirling et al., 2002). Lab conditions can also erode ecologically interesting phenotypic variation (*e.g.* the relationship of male song and aggression in crickets; Hedrick and Bunting, 2014), rendering further examination of phenotypic and genetic correlations meaningless. Finally, heritability is not a stable characteristic of a trait—it is contingent on a population’s history of selection, gene flow, and environmental heterogeneity (Visscher et al., 2008). For these reasons, heritability is best assessed in wild populations (though see Dochtermann et al., 2019).

Today, advances in genomics and bioinformatics provide novel opportunities for the estimation of heritability and other quantitative genetics parameters in the field. Our study—in which we collected phenotypic data and DNA in the field, generated genome-wide relatedness estimates, and related these datasets to one another to estimate heritability and co-heritability of condition and song—demonstrates the feasibility of implementing these methods to study both wild, non-model species and the complex behaviors and traits often neglected in traditional genetics analyses. Finally, our data highlight the potential for using such methods to address longstanding evolutionary problems. Harnessing the potential of these techniques and their applications will expand the role of genetics in evolutionary and ecological studies and allow us to shed the confines of the phenotypic gambit.

## Materials and Methods

### Sampling and data collection

We focused on a population of singing mice inhabiting the San Gerardo de Dota valley of Costa Rica, a high-elevation region in the Talamanca Mountain range (>2200 m) characterized by frequent rainfall and its lush cloud forests, grasslands, and paramo (Fig. 1a). Within this region, we sampled mice at five trapping sites over three years (June – August 2014, 2015, and 2016). Trapping sites were located within 6 km^2^ of one another and occurred at elevations between 2200 m and 2600 m (Figure 3a, Table 1). Two of the sites comprised primary and secondary forests (PC and Robles) while the other two comprised grassy pastures (QERC, MH).

We captured a total of 168 adult mice and brought them to the Quetzal Education and Research Center (QERC) for processing (*N* = 32 females and 134 males). This number was male-biased due to the needs of concurrent projects. We recorded the hindfoot length, anogenital distance, and mass of each focal mouse. Mice were housed singly in 28 × 28 × 28 cm^3^ PVC-coated wire mesh cages and provided with food and water *ad libitum*. Each mouse was given at least one day of acclimation before recording sessions.

Focal animals were moved in their home cage into a 42 × 42 × 39 cm^3^ acoustic isolation chamber the night before recording. Recording sessions took place between 500h and 1100 h in the morning. We have previously described lab methods for obtaining and recording singing mouse songs (Burkhard et al., 2018; Campbell et al., 2010); briefly, we recorded both songs produced in response to playback stimuli and songs spontaneously produced in two hours of silence. Vocalizations were recorded using ACO Pacific microphones on Tucker-Davis RX6 hardware at 32-bit, 195.3 kHz resolution. We visualized the spectrogram of each recording in MATLAB and excluded poor-quality recordings from analysis.

Between 1400 and 1700 h on the day of recording, we euthanized focal animals. Trunk blood was collected within two minutes of sacrifice and immediately centrifuged for plasma extraction. We then harvested fresh liver tissue and counted the number of nematode endoparasites present in the stomach and body cavity. Plasma and liver tissue were stored at −20°C until analysis.

### Song analysis

#### Acoustic measurement

We successfully obtained 467 vocalizations from 111 mice (*N* = 24 females, 87 males). Vocalizations were bimodally distributed, ranging in length from 0.3 s to 9.9 s long (mean ± s.d. = 6.3 ± 1.8 s). We operationally define long songs as those four seconds or longer and retained long songs from 108 mice (*N* = 24 females, 84 males, Table 2). We assessed acoustic properties of each song using custom script in MATLAB. We measured 21 parameters describing changes to spectral, temporal, and amplitude characteristics throughout a song and 5 parameters describing whole-song characteristics (mean bandwidth, song length, note number, entropy, and dominant frequency; Table S1) (Burkhard et al., 2018; Campbell et al., 2010). To these 26 parameters, we added three additional whole-song measures, trill rate (notes/s), and minimum and maximum frequency of song.

#### Multivariate analysis of acoustic variation

To characterize dimensions of acoustic variation, we performed principal components analysis (PCA) on 26 acoustic parameters (Table S1) in R version 3.5.3 (R Core Team, 2016). These variables were selected to accommodate comparison of unpublished data to previously reported data (Burkhard et al., 2018). The first two principal components (PCs) had λ > 4.9 and explained 42.2% of overall variance (Figure 2b). To aid interpretation, we multiplied PC1 scores by −1 (Jolliffe, 2002). Songs scoring strongly and positively on PC1 were longer, had longer beginning notes, and had shorter inter-note intervals than those scoring strongly and negatively on PC1. Songs that loaded strongly and positively on PC2 had a lower dominant frequency, greater mean bandwidth, began at lower frequencies, and ended on higher frequencies than those scoring strongly and negatively on PC2. As these results resembled previously published work (Burkhard et al., 2018), we interpreted PC1 as individual variation in “song effort” and PC2 as variation in “frequency modulation” and retained these as composite acoustic variables for subsequent models (Figure 2c).

#### Repeatability

Because repeatability describes the proportion of total phenotypic variation due to differences between individuals, the metric is often used to estimate the upper limits to a trait’s heritability (Bell et al., 2009; Boake, 1989, though see Dohm, 2002). We estimated the repeatability for each of the eight whole-song measures and for PC1 and PC2 of song. Repeatability is often estimated as the intraclass correlation coefficient (RA., 1954; Wolak et al., 2012), such that:

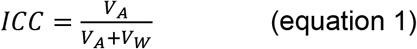

where *V*_*A*_ is the variance among individuals, *V*_*W*_ is the variance within individuals, and total phenotypic variation *V*_*P*_ = *V*_*A*_ + *V*_*W*_. Intraclass correlation coefficients were calculated by fitting mixed-effect models using the *rptR* R-package (v0.9.22, Stoffel et al., 2017) with animal ID as a random effect. Confidence intervals were estimated with parametric bootstrapping for 1000 iterations, and p-values were calculated with likelihood ratio tests.

### Assessment of body condition

#### Morphometric evaluation

Residual body mass (RBM) was determined by regressing mass on skeletal length (*i.e.* hindfoot length) and calculating the residual error in grams for each animal. RBM, a traditional proxy for total energy reserves, was included in subsequent analyses of body condition.

We also measured anogenital distance (AGD), a morphological cue of prenatal exposure to androgens. We performed Z-score normalization on AGD (AGD_z_); because AGD was sexually dimorphic (Welch two sample t-test; *T*= −12.5, *P* < 2.2e-16), it was transformed for each sex separately. Thus, female AGD_z_ > 1.5 corresponds to females with AGD ≥ 5.0 mm whereas male AGD_z_ > 1.5 corresponds to males with AGD ≥ 7.5 mm. Because variation in AGD predicts individual differences in “masculinity” but not body condition itself, we considered AGD as a covariate separate from body condition metrics in subsequent analyses.

#### Plasma assays

Plasma samples collected in 2015 were assayed by the Mouse Metabolic Phenotyping Center at the University of Cincinnati, Ohio. Those collected in 2016 were assayed by the Mouse Metabolic and Phenotypic Core at the Baylor College of Medicine. Only male samples were processed. We selected assays for cholesterol, phospholipids, non-esterified fatty acids (NEFA), glucose, and triglycerides (TG) and the hormones adiponectin, insulin, and leptin. Not every male had enough plasma for all assays, so assays were assigned an order of priority.

Assays for 2015 samples were completed in the following order: 1) glucose, 2) lipid profile (TG, cholesterol, phospholipids, NEFA), and 3) multiplex ELISA assay (insulin, leptin, adiponectin). As results from these first analyses suggested a strong relationship between adiposity hormones and singing behavior (Burkhard et al., 2018), we modified the order for 2016 assays to the following: 1) multiplex ELISA, 2) glucose, and 3) lipid profile. As a result, sample sizes vary across plasma measures (Table 2). Values for plasma measures did not significantly differ between years and were thus pooled for further analysis.

#### Multivariate analysis of variation in body condition

We performed exploratory PCA and assessed a correlation matrix of all body condition parameters (*i.e.* RBM and plasma measures) to examine how body condition varied among individuals. To maximize our statistical power, we ultimately focused on the subset of individuals (*N* = 52) with each of the following parameters: RBM, glucose, leptin, insulin, adiponectin, and triglycerides. We performed PCA on these condition measures in R to create composite variables for downstream analyses. Three PCs had λ > 1.0, explaining 34.5%, 22.2%, and 17.2% of total variance, respectively. Glucose, insulin, and triglycerides strongly loaded on PC1; RBM and adiponectin contributed most to PC2; and leptin and adiponectin contributed strongly and in different directions to PC3 (Table S2). We retained these three components as composite variables in downstream analyses (*i.e.* PC1con, PC2con, PC3con).

### Library preparation and variant calling

We extracted DNA from thawed liver tissue using Qiagen DNeasy Blood and Tissue kits. DNA was first quality-checked by gel electrophoresis and then quantified with a PicoGreen fluorometer quantitator and Quant-iT™ PicoGreen™ dsDNA Assay Kit (Invitrogen™, ThermoFisher Scientific). The Genomic Sequencing and Analysis Facility (GSAF; University of Texas, Austin) performed double-digest restriction site associated DNA sequencing (ddRAD-seq; Peterson et al., 2012) library preparation on normalized DNA concentrations. The restriction enzymes *EcoR1* and *MspI* (New England BioLabs) were selected to shear DNA with Pippen size selection for fragments of 335-435 bp. Each sample was tagged with a unique five-nucleotide-long barcode and then pooled in groups of 24 for sequencing. Samples included biological replicates to check for procedural error. Multiplexed samples were then sequenced on one lane of either the HiSeq 2500 or NextSeq 500 (Illumina, Inc.), depending on availability. Lanes also included samples for studies not described here.

After inspecting the quality of raw reads with FastQC (Andrews, 2010), we demultiplexed and trimmed reads to 125 bps using the *process_radtags* command from the Stacks pipeline (v1.46.0; Rochette and Catchen, 2017). Trimmed reads were mapped to the *S. teguina* reference genome (unpublished) using Bowtie 2 (v2.3.2; Langmead and Salzberg, 2012), and resulting *sam* files were converted to *bam* files using SAMtools (v1.10; Li et al., 2009). We used ANGSD (v0.929-13; Korneliussen et al., 2014) to assess base qualities and coverage depth of resulting bam files, excluding poor-quality and poor-coverage samples from downstream analyses.

Samples that passed these initial filtering steps were then further filtered with ANGSD to retain only loci with minor allele frequency (MAF) above 0.1 and which were present in at least 80% of all individuals. We detected sex-linked markers using VCFTOOLS (v0.1.15; Danecek et al., 2011) and then excluded them with the -rf option in ANGSD. Finally, we used ANGSD to call variants and convert genotype data into various formats, including identity-by-state matrices (ibsmat), mutation annotation format files (maf), variant calling files (vcf), and genetic covariance matrices (covmat).

### Population structure and genomewide relatedness

Pairwise genetic distance between trapping locations (*F*ST) was estimated using VCFTOOLS. We also used the R-package vegan (v.2.5.6; Oksanen et al., 2019) to perform principal coordinates analysis (PCoA) to help resolve genetic distance. Permutational multivariate analysis of genetic distance (PERMANOVA) explained by trapping site was performed with the *adonis* function in vegan.

There are many methods available to calculate genetic relatedness which produce a variety of data formats (Eu-ahsunthornwattana et al., 2014). Certain combinations of data formats with heritability estimation method are more compatible and straightforward than other combinations, however. For use with *GCTA* software (see Inference of SNP-based heritability), we produced a population genomic relationship matrix (GRM) in PLINK (v1.90b6.7; Chang et al., 2015), which uses a method-of-moments approach to estimate the proportion of the genome at which two individuals share 0, 1, or 2 alleles identical by descent (IBD; Purcell et al., 2007). PLINK operates on ‘hard-call’ genotypes and is thus best suited for high-coverage, high-quality reads. To check the accuracy of PLINK relatedness estimation on our data, we also estimated relatedness using NgsRelate software (v2; Korneliussen and Moltke, 2015), which uses a maximum likelihood approach on genotype likelihoods to estimate the proportion of alleles two individuals share IBD. Relatedness estimates from NgsRelate, a method based on genotype likelihoods, were strongly correlated with estimates from PLINK (*R*^*2*^ = 0.90, *P* < 2.2e-16; Fig. S1). Finally, for use with *brms* (see Inference of SNP-based heritability), we produced a population genetic covariance matrix using ANGSD, which uses genotype likelihoods rather than hard-called genotypes.

### Phenotypic correlations between condition and song

To ask how condition phenotypes related to singing behaviors, we fitted mixed models using the R-package *glmmTMB* (v1.0.2.1; Brooks et al., 2017). We focused on the eight whole-song measures, song rate, and the first two song PCs as response phenotypes. Whole-song measures were transformed by Z-score normalization before model fitting. Whole-song measures and song PCs were fitted assuming Gaussian distributions, while song rate was fitted assuming a negative binomial distribution with a logit link function. Each response variable was initially fitted with a full model, which included PC1, PC2, and PC3 of condition and AGD_z_ as covariates. Composite variables, rather than individual parameters, were included to reduce multicollinearity of parameters and to mitigate interpretation of results. Trapping site (Site) and in dividual identity (1|ID) were also included as a fixed factor and a random effect, respectively. Because only male plasma samples were processed, we did not include sex in the models. Finally, we specified the number of days spent in acclimation (Days) as an observation-level dispersion factor to account for data overdispersion (Harrison, 2014). After fitting the full model, we used R-package *buildmer* to perform a backwards stepwise elimination procedure based on the change in AIC (v1.6; Voeten, 2020). We report the estimates, SEs, and p-values for all terms in the final models and the change in AIC between initial and final models. Model results were visualized using the *effects* package (v4.1.4; Fox and Hong, 2009; Fox and Weisberg, 2018), and the *emmeans* (v1.5.0; Lenth, 2020) and *boot* packages were used to help interpret results (v1.3.25; Canty and Ripley, 2020).

### Quantitative genetics

#### Inference of SNP-based heritability

We estimated h2 values for the eight whole-song measures, PC1 and PC2 song, RBM, AGDz, and PC1, PC2, and PC3 condition using animal models implemented with both a maximum likelihood estimation (MLE) approach and a Bayesian Hamiltonian Monte Carlo approach. The MLE method was implemented with *GCTA* standalone software (v1.92.1; Yang et al., 2011), and the Bayesian approach was implemented using the *brms* R-package (v2.13.5; Bürkner, 2018, 2017). While easy to implement and fast to run, GCTA and many related frequentist algorithms cannot accommodate more complex models or non-parametric assumptions (*e.g.* non-normally distributed phenotypes; *e.g.* de Villemereuil, 2019). Conversely, linear (or generalized linear) mixed models (GLMM) implemented with Bayesian approaches are less user-friendly and more computationally intensive but are more robust to scarce data and complex models and are easy to interpret (de Villemereuil, 2019; Morrissey et al., 2014; Villemereuil et al., 2013).

Animal model approaches rely on estimating heritability by correlation between genetic similarity and phenotypic similarity (Wilson et al., 2010).The most basic animal model assumes a normally distributed phenotype, where a trait (*y*) in an individual (*i*) can be described as:

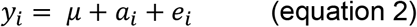

where *μ* is the population mean; *a*_*i*_ is the breeding value, or effect of individual *i*’s genotype on the trait; and *e*_*i*_ is a residual term. This equation is specified as a mixed model containing both fixed and random effects with individual genotype treated as a random effect such that:

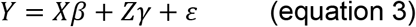

where *Y* is a vector of phenotypic values, *X* is a fixed variable, *β* is the fixed variable effect size, *Z* is the GRM, γ is a vector of random effects, and ε is a residual random effect (non-genetic).

Narrow-sense heritability was calculated as

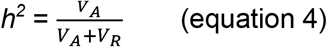

where *V*_*A*_ is additive genetic variance, *V*_*R*_ is residual variance, and *V*_*P*_, is total phenotypic variance, = *V*_*A*_ + *V*_*R*_.

##### GCTA models

Models were fitted with *GCTA* with sex (Sex) and trapping site (Site) included as fixed effects and PLINK-derived GRM as a random effect.

##### Brms models

Heritability estimates for each trait were estimated with models including ANGSD-derived genetic covariance matrix as a random effect, and Sex (excluding models involving only one sex, *i.e.* heritability of PC condition variables) and Site as fixed effects. Where applicable, response variables were log-transformed and scaled to address skew. We specified gaussian family distributions for all variables except song rate, which we fitted assuming a negative binomial distribution. Each model was run with 4 chains of 5000 iterations, with burn-in of 1000, and thin of 1 and provided with weakly informative priors. Model output was checked for proper mixing and convergence by inspection of autocorrelation and diagnostic plots. We took the mean of values from the posterior distribution to calculate heritability and report 95% highest posterior density credible intervals (HPDI) in Table 4. We calculated heritability following equation 4; residual variance for negative binomial models was calculating following (Matos et al., 1997). Credible intervals contain the most probable values (*i.e.* 95%, in this case) of the estimated parameter. Unlike a confidence interval, credible intervals have distributional information such that the most probable values are closest to the point estimate and the width indicates certainty.

### Genetic correlation of song and condition

The proportion of covariance between two traits explained by genetics is calculated by the following equation:

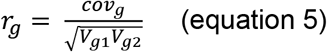

Where *cov*_*g*_ is the genetic covariance, *V*_*g1*_ is the additive genetic variance for trait 1, and *V*_*g2*_ is the additive genetic variance for trait 2. To determine whether traits were genetically correlated, we ran bivariate models in *brms* between all pairwise comparisons of condition and song variables that were found to have statistically clear phenotypic correlations (see Phenotypic correlations between condition and song). These models used the same data and parameters as the univariate models described above for *brms* models. Genetic correlations were considered significant in magnitude if the 95% highest posterior density credible intervals (HPDI) excluded zero.

## Supporting information

Mouse phenotypic data

## Acknowledgments

We thank R. Westwick, K. Garner, W. Betty, and the QERC staff for their assistance in the field. We also thank J. Podnar for support with RAD-seq; B. Goetz and J. Sardell for advice on bioinformatics analysis; P. Vullioud, P. de Villemereuil, and M. Modrák for advice on statistical analysis; G. Otto, P. Saha, and P. Tso for guidance on bloodwork; and the Phelps lab and M. Ryan for comments on drafts of this manuscript.

## Ethics

All animal procedures were approved by the Institutional Animal Care and Use Committee at the University of Texas and the Costa Rican Ministerio de Ambiente y Energía.

## Notes

### Competing Interest Statement

The authors have declared no competing interest.

### Summary of Updates

Abstract and author information updated

