## Supplementary Figures and Tables for "Genomic heritability of song and condition in wild singing mice"

*Table of Contents*

**Figure S1**: Genomic pairwise relatedness in population of singing mice	p 2

**Table S1:** PC unrotated loadings of acoustic measures	p 3

**Table S2:** PC unrotated loadings of condition measures	p 4

**Table S3:** ICC estimates for song	p 5

**Table S4:** GCTA estimates for song and condition	p 6

**Table S5:** Predictors of acoustic parameters in Alston’s singing mice	p 7

**Figure S1:** *(A)* Histogram of pairwise comparisons of genomic relatedness between individual mice. Zoomed-in panel highlights relatedness equivalent to half-sib and full-sib or parent-offspring relatedness. *(B)* Relatedness estimates from ngsRelate are consistent with estimates from plink (*R^2^ = 0.91, P* < 2.2e^-16^).

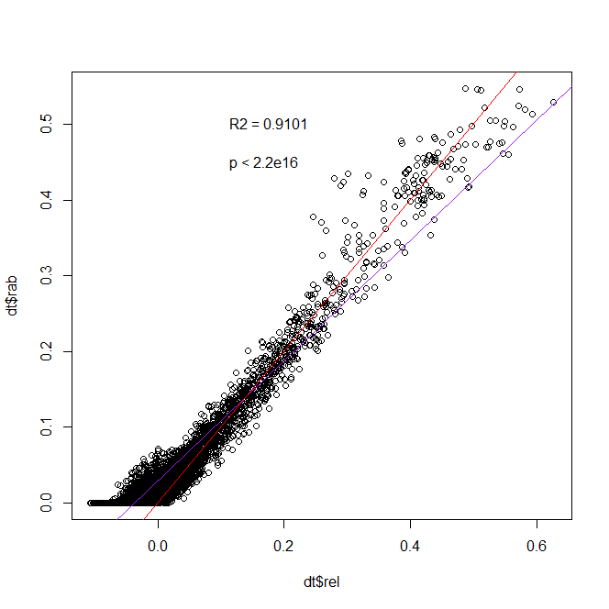

B

Plink relatedness

ngsRelate relatedness

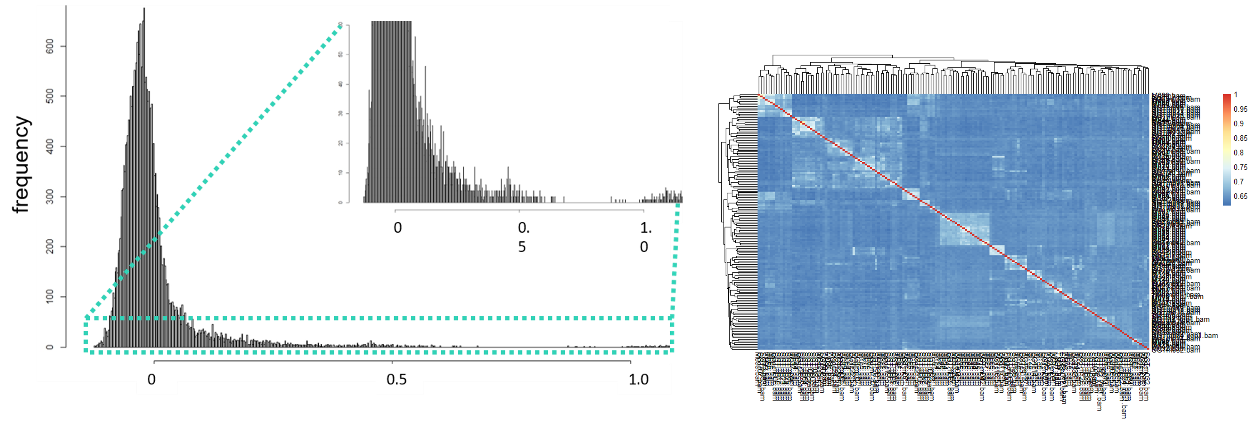

A

**Table S1:** PC song unrotated loadings of acoustic measures. Bold are measures that loaded strongest on varimax rotated components. Italicized are five whole-song descriptors; the others describe changes in properties throughout a song. PC1 has been transformed by multiplying by -1 to aid interpretation.

|  |  | PC1 | PC2 |
| --- | --- | --- | --- |
| *call_length* | LEN | **0.21** | 0.03 |
| *note_num* | NUM | **0.30** | 0.08 |
| *entropy* | ENT | **-0.08** | **-0.21** |
| *call_BW_mean* | BW | 0.02 | **-0.18** |
| *call_DF* | DF | 0.04 | **-0.11** |
| note.durs_a | DURa | 0.05 | -0.08 |
| note.durs_b | DURb | **-0.25** | -0.01 |
| note.durs_c | DURc | **0.17** | 0.06 |
| INI_a | INIa | **-0.32** | -0.06 |
| INI_b | INIb | **0.25** | 0.08 |
| INI_c | INIc | **-0.22** | -0.02 |
| rel.pk.amps_a | PAa | **0.34** | 0.05 |
| rel.pk.amps_b | PAb | **-0.32** | -0.10 |
| rel.pk.amps_c | PAc | -0.05 | 0.13 |
| rel.qpk.amps_a | QAa | **0.33** | 0.06 |
| rel.qpk.amps_b | QAb | **-0.30** | -0.11 |
| rel.qpk.amps_c | QAc | -0.03 | 0.13 |
| FMa_a | FMAa | 0.04 | -0.38 |
| FMa_b | FMAb | -0.06 | 0.40 |
| FMa_c | FMAc | 0.10 | -0.38 |
| FMb_a | FMBa | -0.10 | 0.33 |
| FMb_b | FMBb | 0.07 | -0.38 |
| FMb_c | FMBc | -0.15 | 0.33 |
| FMc_a | FMCa | **0.23** | 0.05 |
| FMc_b | FMCb | **-0.15** | -0.02 |
| FMc_c | FMCc | 0.10 | -0.07 |

**Table S2:** PC condition unrotated loadings. Bolded are terms that loaded strongly on different components, as indicated by varimax rotation.

|  | PC1 | PC2 | PC3 |
| --- | --- | --- | --- |
| RBM | -0.06419 | **0.749767** | -0.15857 |
| Leptin | -0.21574 | 0.267831 | **0.79675** |
| Glucose | **-0.6096** | -0.2851 | -0.27489 |
| Insulin | **-0.47569** | 0.304899 | 0.093295 |
| Triglycerides | **-0.58542** | -0.06263 | -0.14316 |
| Adiponectin | 0.093383 | **0.43353** | **-0.48506** |

**Table S3:** ICC for various acoustic parameters. In italics are 7 whole-song descriptors; other measures describe changes in properties throughout a song. Each measure’s abbreviation corresponds to those plotted in Fig. 2a.

| Parameters | Abbreviation | ICC | 95% CI | | SE |
| --- | --- | --- | --- | --- | --- |
| *trill rate* | *NA* | 0.33 | 0.22 | 0.42 | 0.05 |
| *max Hz* | *NA* | 0.56 | 0.46 | 0.64 | 0.05 |
| *min Hz* | *NA* | 0.76 | 0.69 | 0.81 | 0.03 |
| PC1 of song | *NA* | 0.45 | 0.37 | 0.52 | 0.06 |
| PC2 of song | *NA* | 0.39 | 0.30 | 0.47 | 0.06 |
| *song length* | LEN | 0.33 | 0.25 | 0.43 | 0.05 |
| *note number* | NUM | 0.30 | 0.20 | 0.37 | 0.04 |
| *entropy* | ENT | 0.57 | 0.52 | 0.66 | 0.04 |
| *mean BW* | BW | 0.76 | 0.72 | 0.81 | 0.02 |
| *DF* | DF | 0.43 | 0.31 | 0.49 | 0.05 |
| note.durs_a | DURa | 0.01 | 0.00 | 0.05 | 0.01 |
| note.durs_b | DURb | 0.06 | 0.00 | 0.11 | 0.03 |
| note.durs_c | DURc | 0.34 | 0.24 | 0.42 | 0.05 |
| INI_a | INIa | 0.00 | 0.00 | 0.05 | 0.01 |
| INI_b | INIb | 0.00 | 0.00 | 0.05 | 0.01 |
| INI_c | INIc | 0.02 | 0.00 | 0.07 | 0.02 |
| rel.pk.amps_a | PAa | 0.10 | 0.03 | 0.17 | 0.04 |
| rel.pk.amps_b | PAb | 0.23 | 0.13 | 0.30 | 0.04 |
| rel.pk.amps_c | PAc | 0.36 | 0.27 | 0.43 | 0.04 |
| rel.qpk.amps_a | QAa | 0.13 | 0.06 | 0.20 | 0.04 |
| rel.qpk.amps_b | QAb | 0.24 | 0.15 | 0.31 | 0.04 |
| rel.qpk.amps_c | QAc | 0.39 | 0.29 | 0.47 | 0.05 |
| FMa_a | FMAa | 0.00 | 0.00 | 0.04 | 0.01 |
| FMa_b | FMAb | 0.10 | 0.03 | 0.15 | 0.03 |
| FMa_c | FMAc | 0.35 | 0.28 | 0.46 | 0.05 |
| FMb_a | FMBa | 0.00 | 0.00 | 0.05 | 0.01 |
| FMb_b | FMBb | 0.18 | 0.04 | 0.18 | 0.04 |
| FMb_c | FMBc | 0.52 | 0.38 | 0.55 | 0.04 |
| FMc_a | FMCa | 0.00 | 0.00 | 0.05 | 0.01 |
| FMc_b | FMCb | 0.12 | 0.04 | 0.18 | 0.04 |
| FMc_c | FMCc | 0.61 | 0.51 | 0.66 | 0.04 |

**Table S4:** GCTA estimates. 95% CI = 95% confidence intervals.

| Type | Trait | V_G_/V_P_ | SE | P | 95% CI |
| --- | --- | --- | --- | --- | --- |
| development | AGD | 0.03 | 0.16 | 4.12E-01 | -0.28-0.35 |
| condition | PC1con | 0.96 | 0.59 | 3.79E-01 | -0.20-2.11 |
| condition | PC2con | 0.03 | 0.34 | 4.63E-01 | -0.64-0.70 |
| condition | PC3con | 0.21 | 0.49 | 3.42E-01 | -0.74-1.16 |
| condition | RBM | 0.14 | 0.17 | 1.58E-01 | -0.19-0.48 |
| condition | Hindfoot | 0.22 | 0.24 | 2.20E-01 | -0.25-0.70 |
| song | Entropy | 0.17 | 0.31 | 2.98E-01 | -0.43-0.77 |
| song | Max Hz | 0.73 | 0.26 | 9.17E-05 | 0.23-1.23 |
| song | Mean BW | 1.00 | 0.27 | 5.53E-06 | 0.48-1.52 |
| song | Min Hz | 0.91 | 0.24 | 3.64E-06 | 0.43-1.39 |
| song | Note num | 0.00 | 0.28 | 5.00E-01 | -0.54-0.54 |
| song | PC1 song effort | 0.12 | 0.29 | 3.38E-01 | -0.45-0.69 |
| song | PC2 freq mod | 0.63 | 0.33 | 1.28E-02 | -0.02-1.29 |
| song | Song DF | 0.35 | 0.30 | 7.26E-02 | -0.23-0.94 |
| song | Song length | 0.00 | 0.26 | 5.00E-01 | -0.51-0.51 |
| song | Song rate | 0.61 | 0.31 | 2.02E-02 | 0.01-1.22 |
| song | Trill rate | 0.00 | 0.27 | 5.00E-01 | -0.52-0.52 |
| song | Fma_a | 0.01 | 0.23 | 4.78E-01 | -0.43-0.46 |
| song | Fma_b | 0.16 | 0.29 | 2.83E-01 | -0.41-0.73 |
| song | Fma_c | 0.46 | 0.35 | 1.41E-01 | -0.24-1.15 |
| song | Fmb_a | 0.00 | 0.21 | 5.00E-01 | -0.41=0.41 |
| song | Fmb_b | 0.05 | 0.19 | 3.72E-01 | -0.32-0.42 |
| song | Fmb_c | 0.24 | 0.31 | 2.15E-01 | -0.37-0.85 |

**Table S5:** Predictors of acoustic parameters in Alston’s singing mice. We report coefficients (estimate), SEs, and P values from GLMMs. Bold terms were included in the final model. Terms in italics were included in the initial model and dropped during model specification and displayed with estimates and probabilities from the initial model. AGD = anogenital distance; disp = dispersion parameter.

| Predictors | Δ AIC | estimate | SE | P value |
| --- | --- | --- | --- | --- |
| Entropy | 7.6 |  |  |  |
| PC1con |  | *0.04* | *0.07* | *0.56* |
| **PC2con** |  | **-0.18** | **0.09** | **0.04** |
| **PC3con** |  | **0.16** | **0.10** | **0.11** |
| AGD.c |  | *0.03* | *0.17* | *0.85* |
| **disp=~Days.in.lab** |  | **-0.09** | **0.25** | **0.71** |
| Song length | 13.9 |  |  |  |
| PC1con |  | *-0.04* | *0.08* | *0.66* |
| **PC2con** |  | **0.32** | **0.10** | **0.00** |
| PC3con |  | *-0.01* | *0.11* | *0.96* |
| AGD.c |  | *0.28* | *0.21* | *0.18* |
| **disp=~Days.in.lab** |  | **-1.83** | **0.89** | **0.04** |
| Note number | 13.6 |  |  |  |
| PC1con |  | *-0.06* | *0.08* | *0.44* |
| **PC2con** |  | **0.32** | **0.10** | **0.00** |
| PC3con |  | *-0.05* | *0.10* | *0.64* |
| AGD.c |  | *0.31* | *0.21* | *0.14* |
| **disp=~Days.in.lab** |  | **-1.77** | **0.91** | **0.05** |
| Trill rate | 9.5 |  |  |  |
| PC1con |  | *-0.01* | *0.09* | *0.91* |
| PC2con |  | *-0.04* | *0.12* | *0.71* |
| **PC3con** |  | **-0.11** | **0.11** | **0.34** |
| AGD.c |  | *-0.10* | *0.25* | *0.70* |
| **disp=~Days.in.lab** |  | **-0.78** | **0.27** | **0.00** |
| PC1 song | 8.1 |  |  |  |
| **PC1con** |  | **0.45** | **0.20** | **0.82** |
| PC2con |  | *-0.10* | *0.19* | *0.59* |
| PC3con |  | *-0.09* | *0.24* | *0.71* |
| **AGD.c** |  | **0.76** | **0.36** | **0.04** |
| **disp=~Days.in.lab** |  | **-0.84** | **0.28** | **0.00** |
| Song rate | 11.2 |  |  |  |
| PC1con |  | *0.00* | *0.09* | *0.96* |
| PC2con |  | *-0.10* | *0.12* | *0.39* |
| **PC3con** |  | **0.21** | **0.10** | **0.04** |
| AGD.c |  | *0.06* | *0.24* | *0.82* |
| **disp=~Days.in.lab** |  | **1.90** | **0.80** | **0.02** |
| PC2 song | 16.3 |  |  |  |
| **PC1con** |  | **0.23** | **0.16** | **0.14** |
| PC2con |  | *0.10* | *0.23* | *0.67* |
| PC3con |  | *0.22* | *0.21* | *0.31* |
| AGD.c |  | *0.31* | *0.39* | *0.43* |
| **disp=~Days.in.lab** |  | **-0.20** | **0.31** | **0.51** |
| Dominant frequency | 13.7 |  |  |  |
| PC1con |  | *-0.04* | *0.10* | *0.73* |
| PC2con |  | *0.06* | *0.14* | *0.69* |
| **PC3con** |  | **-0.18** | **0.15** | **0.22** |
| AGD.c |  | *0.17* | *0.25* | *0.51* |
| **disp=~Days.in.lab** |  | **-0.28** | **0.31** | **0.38** |
| Min Hz | 11.9 |  |  |  |
| PC1con |  | *0.00* | *0.01* | *0.91* |
| PC2con |  | *0.00* | *0.01* | *0.64* |
| **PC3con** |  | **-0.04** | **0.01** | **0.00** |
| **AGD.c** |  | **-0.02** | **0.01** | **0.14** |
| **disp=~Days.in.lab** |  | **0.36** | **0.29** | **0.21** |
| Max Hz | 6.4 |  |  |  |
| PC1con |  | *0.01* | *0.08* | *0.88* |
| PC2con |  | *-0.01* | *0.11* | *0.95* |
| **PC3con** |  | **0.07** | **0.13** | **0.01** |
| **AGD.c** |  | **0.46** | **0.17** | **0.57** |
| **disp=~Days.in.lab** |  | **0.09** | **0.29** | **0.75** |
| Mean bandwidth | 8.6 |  |  |  |
| **PC1con** |  | **-0.05** | **0.05** | **0.33** |
| PC2con |  | *-0.07* | *0.07* | *0.36* |
| PC3con |  | *-0.02* | *0.07* | *0.80* |
| AGD.c |  | *0.14* | *0.15* | *0.35* |
| **disp=~Days.in.lab** |  | **-1.13** | **0.51** | **0.03** |
